# FreeTrace enables fractional Brownian motion-based single-molecule tracking and robust anomalous diffusion analysis

**DOI:** 10.64898/2026.01.08.698486

**Authors:** Junwoo Park, Nataliya Sokolovska, Clément Cabriel, Asaki Kobayashi, Enora Corsin, Fabiola Garcia-Fernandez, Ignacio Izeddin, Judith Miné-Hattab

## Abstract

Single-molecule tracking (SMT) in live cells reveals how biomolecules explore crowded intracellular environments, yet most tracking software assumes Brownian motion, an approximation that fails when anomalous diffusion dominates. This leads to biased trajectory reconstruction and loss of biophysical information, particularly for the short trajectories typical of intracellular experiments. We present FreeTrace, an SMT framework that reconstructs trajectories under fractional Brownian motion (fBm), incorporating temporal correlations directly into linking with minimal input parameters. A deep neural network estimates diffusion properties (Hurst exponent *H* and generalised diffusion coefficient *K*) for individual trajectories, while an analytical ensemble estimator accurately recovers *H* from trajectories as short as three frames, conditions where mean-squared displacement methods fail. Benchmarking on simulated data demonstrates superior performance across motion types and densities. Applications to chromatin-bound histones, DNA repair proteins in *S*.*c*. yeast and human cells reveal biologically meaningful diffusion subpopulations, with H values consistent with polymer models and confined motion. FreeTrace bridges theoretical anomalous diffusion models and routine biological experiments.

## Introduction

Single-molecule tracking (SMT) has become a central tool for visualising how biomolecules explore their local environment inside living cells [1, 2, 3, 4, 5, 6]. By following individual proteins, nucleic acids, or chromatin *loci*, SMT enables direct measurements of binding dynamics, chromatin organisation, intracellular transport, and phase-separated condensates [7, 8, 9]. State-of-the-art microscopy experiments now routinely generate thousands of trajectories per experiment, yet extracting quantitative biophysical information from these data remains challenging. Most widely used tracking methods are built on the assumption that molecules diffuse as Brownian particles [3, 10, 11]. This approximation simplifies linking detections into trajectories, but it rarely reflects the crowded, heterogeneous, and structured nature of the intracellular environment [12, 13, 14, 15]. Inside nuclei and cytoplasm, molecular motion is strongly influenced by chromatin compaction, macromolecular crowding, transient binding, and active processes [16, 17, 18]. These factors often result in anomalous diffusion, where molecular displacements are temporally correlated, and the mean squared displacement (MSD) no longer grows linearly with time [19, 14, 20, 4]. Ignoring these correlations leads to three critical failures: biased trajectory reconstruction due to incorrect linking probabilities, inaccurate estimates of diffusion coefficients that miss the temporal structure of motion, and loss of biological information about the memory-like properties of molecular exploration [18, 4, 21, 22]. These issues become particularly severe when trajectories are short, as is typical in live-cell imaging [14, 23].

### Fractional Brownian motion as a unifying framework

Biophysical studies have long used fractional Brownian motion (fBm) [24, 25, 26, 27, 25, 28, 20, 29] to describe anomalous diffusion, because it encompasses subdiffusion, Brownian motion, and superdiffusion within a single mathematical framework [30, 31, 32]. In fBm, the MSD scales as ⟨*r*^2^(*t*)⟩ = 2*Kt*^2*H*^, where *K* is the generalised diffusion coefficient and *H* is the Hurst exponent [30, 33, 32, 4]. The Hurst exponent characterises the type of motion: *H* = 0.5 corresponds to Brownian motion, *H <* 0.5 indicates subdiffusion with anticorrelated steps where molecules tend to reverse direction, and *H >* 0.5 indicates superdiffusion with correlated steps where molecules persist in their direction of motion. These correlations (memory-like) arise from the viscoelastic properties of the cytoplasm, transient binding interactions, or active transport processes [16, 17, 18]. Yet, despite its biological relevance, fBm has remained largely inaccessible to the broader SMT community. Existing tracking tools do not incorporate temporal correlations into the linking step and rely on MSD or jump-distance analysis that perform poorly on short trajectories [34, 19]. The difficulty stems from three technical barriers: fBm is computationally expensive to simulate for realistic experimental conditions, analytical estimation of *H* requires long trajectories with many steps, and trajectory linking algorithms optimised for Brownian motion produce systematic errors when applied to anomalous diffusion [10, 11]. As a result, biologists often face a dilemma: use low-density acquisitions to preserve tracking accuracy at the cost of throughput, or use high-density imaging that yields more trajectories but with reduced reliability.

### FreeTrace: Making fBm practical for biological SMT

Here, we introduce FreeTrace, a general SMT analysis framework designed to make fBm-based trajectory reconstruction and diffusion quantification practical for biological experiments. FreeTrace requires only 2 input parameters for molecule detection (detection threshold and point spread function size) and uses a unified tracking algorithm that naturally accommodates subdiffusive, Brownian, and superdiffusive motions without parameter tuning. To address the challenge posed by the short trajectories typical of SMT, FreeTrace integrates two complementary strategies, providing the predicted molecular trajectories from microscopy videos. A deep neural network provides the estimates of diffusion properties (*H* and *K*) for individual trajectories, revealing qualitative trends and enabling population stratification even when individual estimates are biased. In parallel, an analytical ensemble-level estimator uses the distribution of 1D displacement ratios (**Supplementary Note 1**) to accurately infer the Hurst exponent, even when trajectories contain only three frames. This estimator overcomes key limitations of MSD-based analysis by eliminating the need to choose a specific time lag and robustly predicts the expected evolution of molecular motion under fBm. We benchmark FreeTrace on simulated datasets covering a range of motion types (subdiffusive, Brownian, superdiffusive) and molecular densities (5-20 molecules per frame). Across all conditions, FreeTrace reconstructs trajectories and preserves underlying diffusion structure more reliably than widely used SMT software, as quantified by dynamic time warping (DTW) distances for displacement distributions, mean jump distances, angular distributions, and trajectory lengths. We then apply the method to three experimental systems that illustrate distinct biological diffusion regimes: chromatin-bound histones H2B in human osteosarcoma cells, revealing subdiffusion consistent with polymer models of chromatin (*H* ≈ 0.29); the DNA repair factor Rad51 in *S*.*c*. yeast nuclei, showing bounded Brownian motion (*H* ≈ 0.47); and the multi-functional protein human fused in sarcoma (FUS) in human nuclei, revealing two distinct mobility states with different anomalous diffusion properties. These applications illustrate how integrating fBm into trajectory reconstruction provides more accurate and biologically interpretable measurements of molecular motion, bridging the gap between theoretical models of anomalous diffusion and routine biological SMT experiments. FreeTrace participated in the international SPT challenge AnDi [33] in 2024 and ranked first in the video task.

### Overview of the FreeTrace Framework

#### Minimal-parameter localisation

The first strength of FreeTrace lies in its small number of input parameters. Indeed, the parameter-dependent nature of algorithms hinders users’ choice and produces highly parameter-dependent results. Our proposed SMT software, therefore, aims to be less parameterised for reconstructing individual molecular trajectories from a video, while also increasing generality to handle various motion types. FreeTrace requires two input parameters (detection threshold, Gaussian point spread function radius) to control the quantity and quality of detected molecules, and one optional parameter (maximum jump distance) for the reconstruction of trajectories. This optional parameter can be automatically estimated using a sampling method, provided that the number of detected molecules in the video is sufficiently high and the trajectory lengths are long enough for reliable estimation.

The decision of the presence of a molecule is made with the comparison of likelihood ratios [3] (**Supplementary Fig. 1a**, equation 1) between the predefined symmetrical point spread function (PSF) with radius *r* and background signals inside square sliding windows under the Akaike information criterion (AIC). The localisation precision of molecules in SMT is highly dependent on their diffusivity, as molecular motion distorts the PSF in videos, unless molecules are immobile. To extend the generality of localisation, we fit the observed PSF with a bivariate Gaussian function, estimating not only the centres and intensity, but also the xy variances of PSF and the correlation coefficient of the variances (**Supplementary Fig. 1ab**, equation 2). The only two mandatory input parameters of FreeTrace are utilised in this localisation process: the radius *r* of the expected PSF (trivial to estimate and less subject to user bias), and the detection threshold *λ* to control the number of observed particles in a video.

#### Trajectory reconstruction under fBm

The second strength of FreeTrace resides in the generality of its tracking algorithm, in which the reconnection mechanism has been developed beyond the conventional limits of Brownian motion. FreeTrace utilises fractional Brownian motion (fBm) for the reconstruction of molecular trajectories. The trajectory reconstruction under fBm enabled us to increase the generality of SMT software to handle various molecular diffusion types (**Fig. 1ab**) in living cells, including subdiffusive and directed motions, without requiring any changes of input parameters or algorithms. This was possible with the estimation of the Hurst exponent *H* of fBm with deep neural networks for short trajectories. This enables the trajectory for each molecule to be reconstructed considering its motion type in the past, which can be beneficial in heterogeneous environments. Of note, existing tracking methods assume that molecular motion is Brownian, implying that the mechanism is based on a nearest-neighbour approach, as well as additional terms such as the number of detected photons over time for the molecules.

**Fig. 1:**
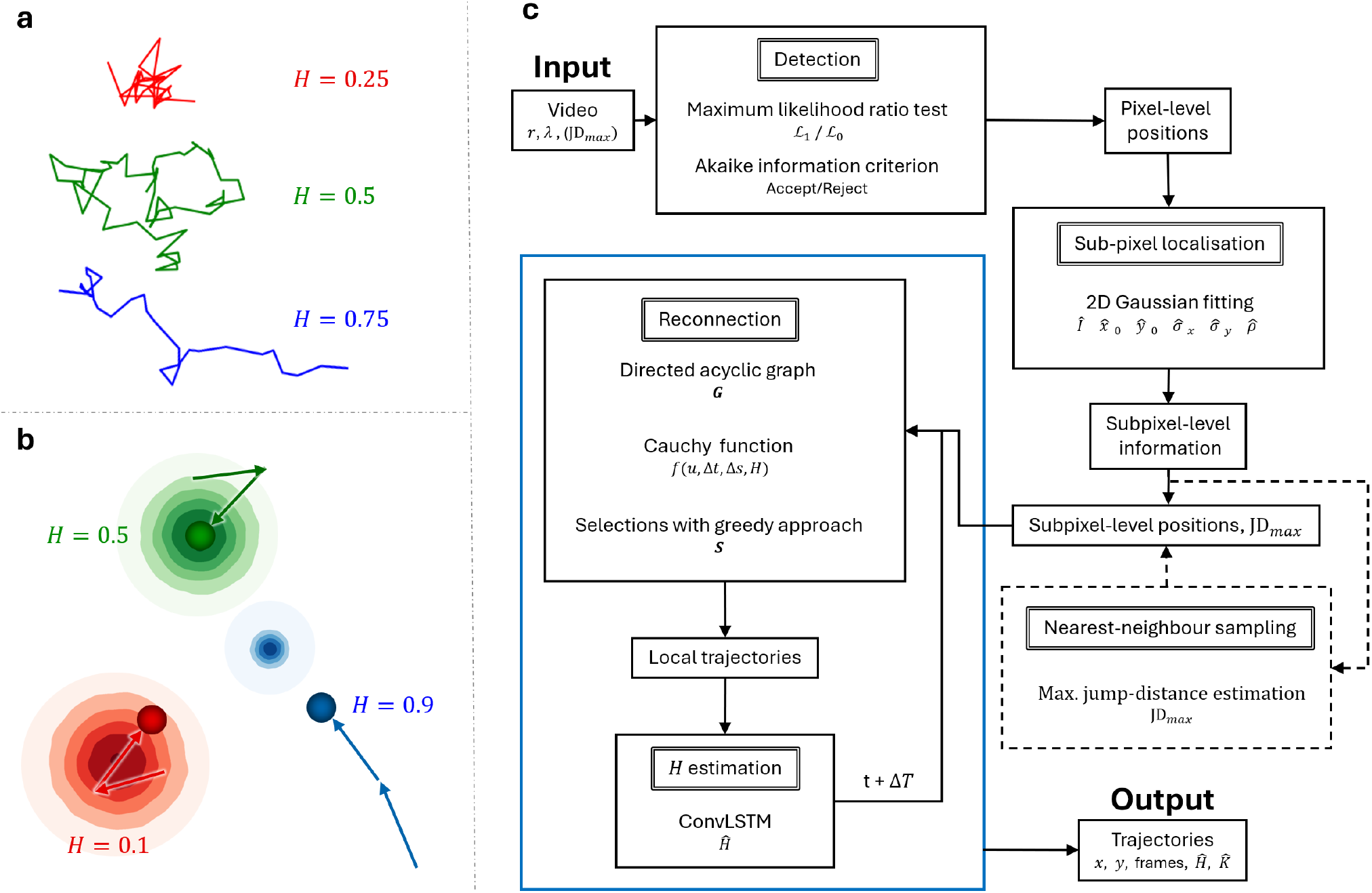
Exemplary fBm trajectories, maximised likelihood ratio test and schematic overview of FreeTrace algorithms. **a**, Examples of different diffusion motions of 2-dimensional fBm trajectories. FBm trajectories can be categorised into subdiffusive motion (red), where *H* ∈]0, 0.5[, superdiffusive motion (blue), where *H* ∈]0.5, 1.0[, and Brownian motion (green), where *H* = 0.5. **b**, The illustration of the probability function implemented in FreeTrace to reconnect the detected molecules with respect to *H*, under the assumption of fractional Brownian motion. The Brownian (green), subdiffusive (red), and superdiffusive (blue) molecules. The most probable next positions of molecules from given *H* are shown as contour maps. The higher probability is represented with a darker colour in the contour map. The estimation of *H* for an individual trajectory is performed with a neural network. **c**, The schematic overview of FreeTrace. The corresponding strategies for each step are shown with output values. The blue box indicates the recursive process over video frames. The dashed-line box indicates a process which can be skipped depending on the input parameter JD_*max*_. FreeTrace accepts two input parameters *r* (PSF radius), *λ* (detection threshold), and one optional parameter JD_*max*_. If JD_*max*_ is not given as an input parameter, an additional process to estimate the maximum jump distance from a given sample is performed. We estimate *H* (Hurst exponent) from the reconstructed past trajectories, and this memory is passed to the next *t* + Δ*T* frame. A DAG (directed acyclic graph) is constructed for Δ*T* frames (**Supplementary Fig. 1c**), and each edge is weighted with the density of the Cauchy function. The most probable paths from *t* to *t* + Δ*T* are selected with a greedy approach. At the end of the video, the DAG containing the reconstructed trajectories is returned.

#### Two-level diffusion estimation

We also present methods for estimating *H* at the two different levels, individual and ensemble level, for single-regime trajectories. The *K* estimation has been explored thoroughly in the literature, considering the localisation errors. However, the estimation of *H* has not been studied extensively in SMT, and *H* is typically inferred by fitting the MSD at the ensemble level. FreeTrace provides the *H* estimate for each individual trajectory using a pre-trained neural network. This estimation at the individual level can reveal trends of molecular behaviours; however, the estimated value can be highly biased when the trajectory is short. To obtain a reliable *H* value from short trajectories at the ensemble level, we present a novel estimation, the Cauchy fitting, which uses the ratio of 1D displacements of trajectories under the assumption of fBm. The estimated *H* explains the memory-like diffusion behaviour of molecules and can be used to interpret biophysical phenomena of molecules in living cells.

## Benchmarking FreeTrace on Simulated Data

We validated the quality of FreeTrace against other publicly available SMT software from five variables: 2D displacement, mean jump-distance, angle, 1D ratio, and trajectory-length distributions (**Table 1, Fig. 2**). If the software has common parameters, we utilised the same values. For the numerical quantification of the trajectory prediction results between the simulated ground-truth (GT) and each software, the dynamic time warping (DTW) [35] distance is utilised to account not only for the shape of the distribution but also for the number of predicted trajectories. The illustration of variables and DTW for the quantification of performances are introduced in **Supplementary Fig. 2**. The DTW returns the optimal match between two sequences, whether of equal or unequal length, and measures the minimum distance between them. The DTW is conceptually similar to the Needleman–Wunsch algorithm [36], which is used for aligning protein or nucleotide sequences. We measured DTW distances on the density function and the cumulative distribution, normalised to the GT. The utilised parameters of each software, visualised DTW paths with the distance score, and the details of each variable are available in **Supplementary Note 1**.

**Table 1:** DTW distances of each software compared to the GT. DTW distance indicates the relative distances of distributions to the GT, where the distributions are normalised to the GT before comparison. The table presents numerical quantifications of the results shown in **Fig. 2** at low and high densities. The distances are obtained with the distance of the density function multiplied by the distance of the cumulative distribution. The DTW distance decreases to zero as the prediction results better match the GT. The values present the rank with DTW distance in parentheses. The number beside the software name indicates the number of parameters and algorithms to produce the results. The parameters of each software to produce results below are available in **Supplementary Table 1**.

|  | FreeTrace (3)<sup>1</sup> | TrackMate (7) | TrackIt (5) | NN<sup>4</sup> (5) |
| --- | --- | --- | --- | --- |
| Density | low \| high |  |  |  |
| 2D disp<sup>2</sup> | 1 (8.6e-3) \| 1 (2.4e-2) | 2 (3.0e-2) \| 2 (4.3e-2) | 4 (1.8e-1) \| 4 (6.8e-1) | 3 (4.5e-2) \| 3 (7.0e-2) |
| MJD<sup>3</sup> | 1 (7.9e-3) \| 1 (1.2e-2) | 3 (6.5e-2) \| 2 (4.5e-2) | 4 (7.9e-2) \| 4 (4.0e-1) | 2 (2.6e-2) \| 3 (4.9e-2) |
| Angle | 1 (2.7e-3) \| 1 (7.8e-3) | 2 (1.2e-2) \| 2 (1.8e-2) | 4 (3.3e-1) \| 4 (3.1e-1) | 3 (1.3e-2) \| 3 (2.2e-2) |
| Length | 2 (1.5e-1) \| 1 (1.8e-1) | 4 (3.3e-1) \| 4 (4.4e-1) | 1 (9.4e-2) \| 3 (3.3e-1) | 3 (1.9e-1) \| 2 (2.6e-1) |
| 1D ratio | 1 (8.3e-3) \| 1 (1.4e-2) | 3 (5.5e-2) \| 3 (7.7e-2) | 4 (2.0e-1) \| 4 (7.2e-1) | 2 (4.3e-2) \| 2 (6.6e-2) |

Rank
<sup>1</sup>Number of input parameters used
<sup>2</sup>Displacement
<sup>3</sup>Mean jump-distance
<sup>4</sup>Nearest-neighbour

**Fig. 2:**
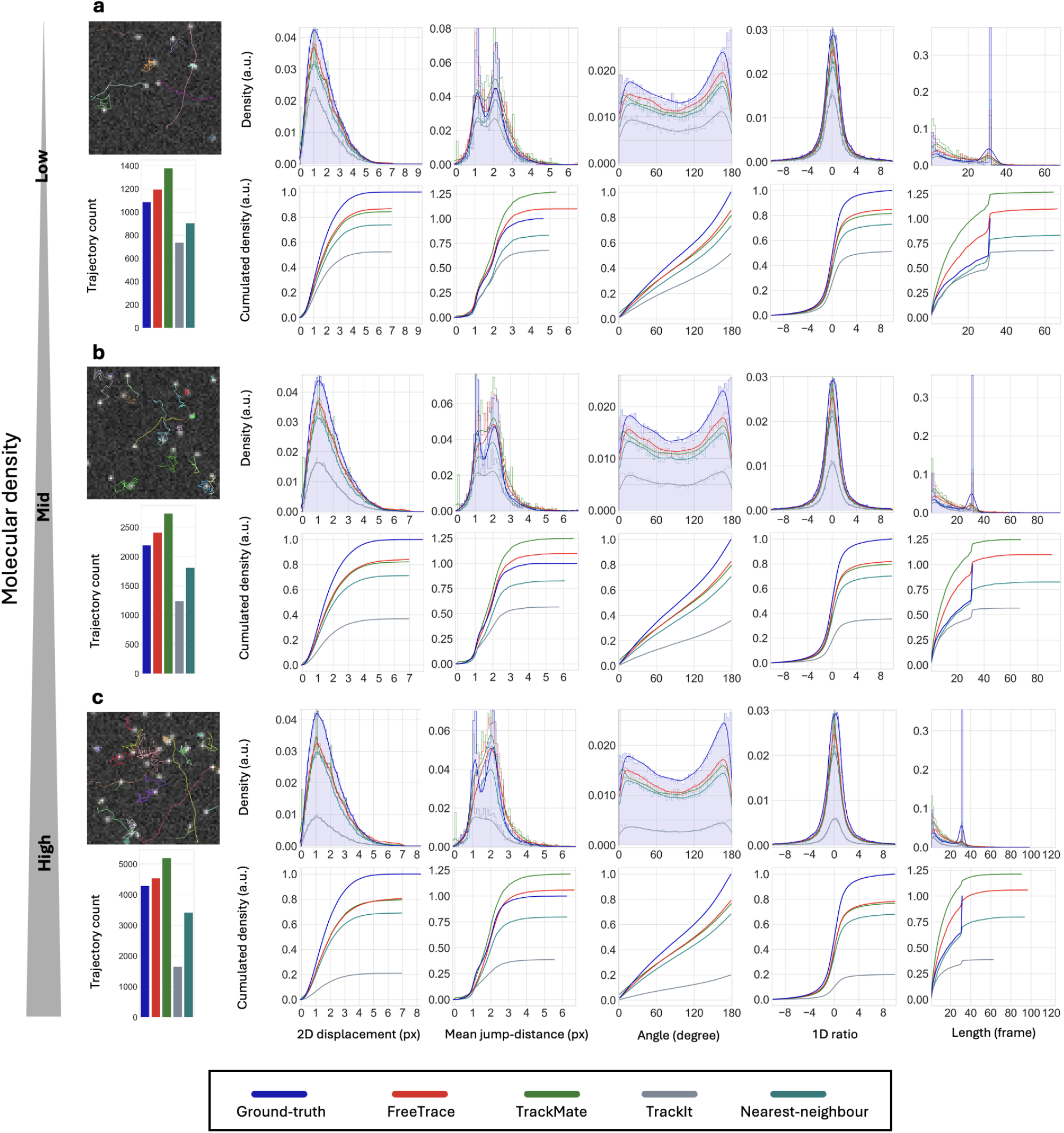
Performance comparison of FreeTrace and SMT software with 5 different distributions on the simulated fBm trajectories. **a**,**b**,**c**, The prediction results at low, mid and high molecular densities, respectively. The average SNR (Signal-to-noise ratio) of the fluorescence signal is 3 in the simulations. The top leftmost figures of **a, b**, and **c** display snapshots of GT trajectories at an arbitrary frame, with the bar chart of the predicted number of trajectories for each software below. We validated the performance of each software with five different variables. The histograms display the density function of each predicted variable, with the curves estimated using a Gaussian kernel. The cumulative distributions of each density function are shown below. Note that the density and distribution functions are normalised to the GT. The closer to the curve of GT, the better the prediction result is in terms of both probability distribution and the number of trajectories. FreeTrace (red) shows a relatively higher accordance with GT (blue) in almost all scenarios, except for the trajectory length at low and mid densities. The numerical quantification of the comparison results utilising DTW (Dynamic Time Warping) is given in **Table 1**. The higher the molecular density, the less accurate the predicted mean jump-distance distribution becomes, losing two peaks due to mislinkage during the trajectory reconstruction step. The explanation of each variable and the parameters of each software are available in the supplementary note. Three types of trajectory motion are simulated with (*K, H*) ∈ { (0.4, 0.1), (1.5, 0.5), (1.5, 0.9)}, *i*.*e*., subdiffusive, Brownian and superdiffusive motion, respectively. The results only contain the predicted trajectories where the length is greater than two. The maximum length of the simulated trajectories in the scenarios is limited to 32.

Three different types of co-existing motions are employed in the simulated videos: i) superdiffusive, ii) Brownian motions, each with the same generalised diffusion coefficient *K*, and iii) subdiffusive motion (**Fig. 2**). Additional scenarios with varying trajectory lengths at high molecular density are also presented in **Supplementary Fig. 4**. Each scenario contains 200 frames from 20 videos, with a 64 × 64 pixel size. Three different molecular densities, 5, 10 and 20 molecules per frame on average, are simulated with three types of trajectory motion with (*K,H*) ∈ {(0.4, 0.1), (1.5, 0.5), (1.5, 0.9)}, subdiffusive, Brownian and superdiffusive motion, respectively. The visualised GT trajectories of each type and a snapshot of the video are shown in the top left-most images in **Fig. 2**. The simulated length of trajectories decays exponentially, with a maximum bounded to 32. We did not simulate motion-changing trajectories to better observe the erroneous linkage at the level of individual trajectories, as it can be observed in the mean jump-distance distribution. The GT of mean jump-distances shows two peaks (**Fig. 2**, 3rd column), as the simulation contains two different *K* values: 0.4 and 1.5. These two peaks are not clearly observable in the displacement distribution due to the small difference in *K* between the motion types. Suppose mis-reconnection (*i*.*e*., erroneous linkage at the trajectory reconstruction step) occurs at the level of individual trajectories between *K* values of 0.4 and 1.5. In that case, the two peaks disappear since the mean jump-distance lies between these two values.

FreeTrace demonstrated its performance in all aspects (**Table 1**) for predicting individual molecular trajectories from video, in accordance with the GT in five variables. We can observe that mis-reconstruction of molecular trajectory occurred more frequently in high-density videos, as the two peaks in mean jump distance became less distinct in all software. This effect reminds us of the importance of accounting for different perspectives on various motion types, which is largely obscured when we consider only the distribution of 2D displacements. Obviously, the molecular density of videos is crucial for identifying the time point at which molecular motion changes at the individual trajectory level. If the density is high, many predicted time points may be false positives. Therefore, the molecular density should be adjusted at the video acquisition step in real data before performing trajectory prediction.

## Accurate Ensemble Estimation of Anomalous Diffusion

The analysis of molecular diffusivity in fBm heavily relies on the accuracy of *H* estimation, particularly for short trajectories in experimental data. The recent international AnDi challenge [33] demonstrated the potential of AI models for estimating *H* at both the individual and ensemble levels for molecular trajectories, thereby deciphering biological phenomena in cells using SMT. Interestingly, the *H* estimation at the ensemble level is performed using clustering-based methods in the afore-mentioned paper, such as the Gaussian mixture model (GMM), based on the estimated *H* values of individual trajectories. However, the clustering on the estimated *H* contains a very high risk of bias, which is induced by the trajectory predictor and the diffusion property estimators, especially for short trajectories [37]. This becomes more critical when we consider that many molecular trajectories in the nucleus are short with high diffusivity in experimental data. It is natural that the inaccuracy of *H* estimation for individual short trajectories is, of course, inevitable due to the low number of observations. In this section, we present the result of *H* estimation of FreeTrace at both the individual and ensemble levels from the short fBm-simulated trajectories.

**Fig. 3a,b** shows the predicted trajectories with FreeTrace on simulated videos with two populations, the clusters of estimated *H* and *K* from individual trajectories, and the histogram of trajectory length. The left-most images of **Fig. 3a,b** are zooms on predicted trajectories with FreeTrace. **Fig. 3a** consists of two equal-rate populations, subdiffusive and Brownian molecules with *K* and *H* equal to 0.4 and 0.1 for the subdiffusive molecules, and 1.5 and 0.5 for Brownian molecules. **Fig. 3b** also consists of two equal-rate populations, Brownian and superdiffusive molecules with *K* and *H* equal to 1.5 and 0.5 for the Brownian molecules and 1.5 and 0.99 for superdiffusive (directed) molecules. We can observe that FreeTrace identifies two populations forming clusters, as is observable in both visualised trajectories. This proves the potential of AI models to predict the diffusion types of molecules even for short trajectories. However, we can also observe that the estimated centres of clusters (pink cross) are biased from the theoretical *H* (green dot). This clearly shows the risk of bias in the *H* estimate, mainly due to the short length of the trajectories.

**Fig. 3:**
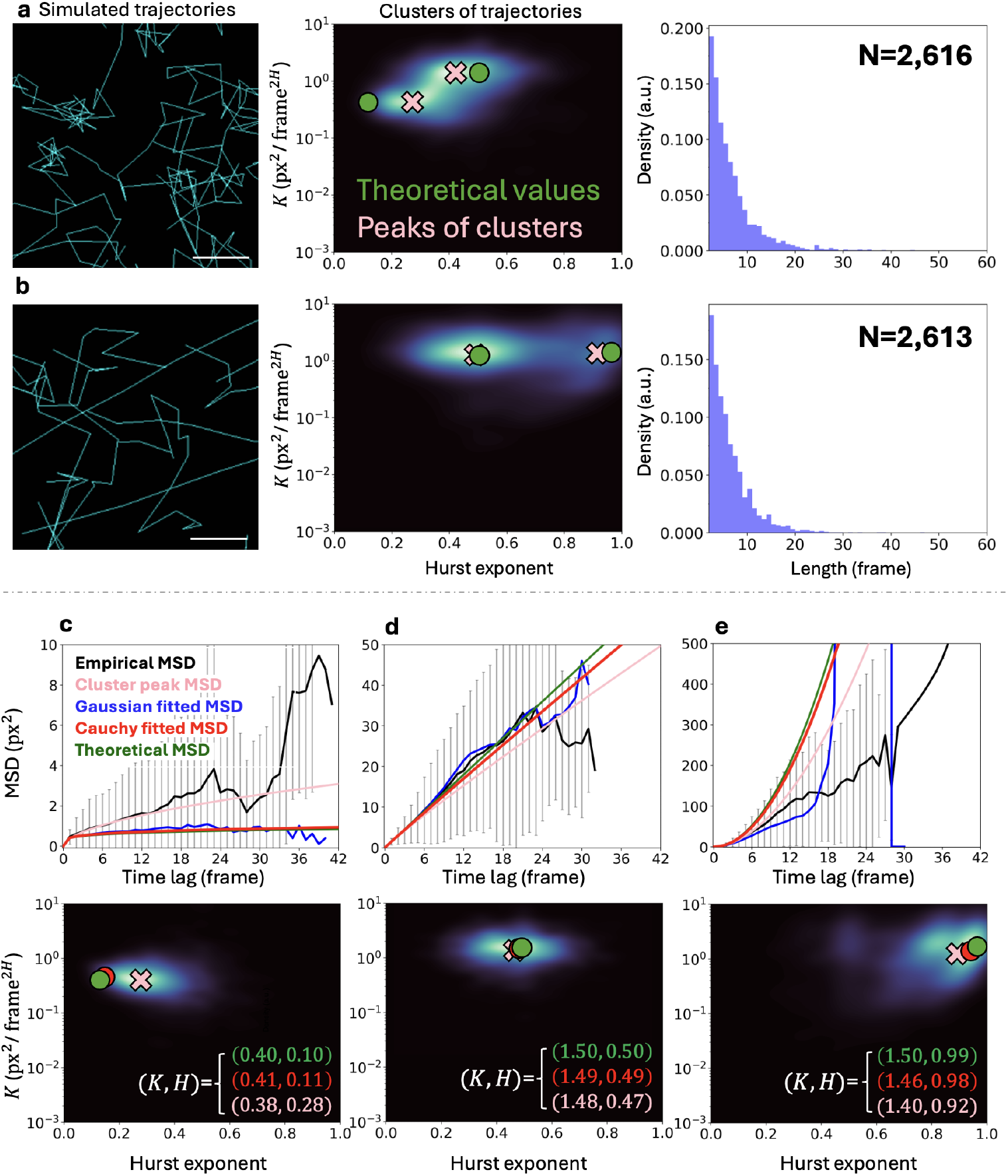
Estimated clusters of *H* and *K* at the individual level predicted with FreeTrace, and an improved *H* estimation for short trajectories at the ensemble level. **a**,**b**, The mixture of predicted fBm trajectories with FreeTrace (left), the clusters identified from the estimated diffusion properties *H* (Hurst exponent) and *K* (generalised diffusion coefficient) of individual trajectories (centre), and the length distribution of the predicted individual trajectories (right). The trajectories exiting the FOV (field of view) are truncated. The result of fBm trajectories in different simulated scenarios: **a** (*K, H*) ∈ {(0.4, 0.1), (1.5, 0.5)} (subdiffusive and Brownian molecules) and **b** with (*K, H*) ∈ {(1.5, 0.5), (1.5, 0.99)} (Brownian and superdiffusive molecules). The green dot indicates the theoretical values of *H* and *K*, and the pink dot indicates the estimated centre of the cluster from individual trajectories. The estimated *H* centres of clusters (pink cross) are biased from theoretical values (green dot) due to the short length of trajectories. **c**,**d**,**e**, The comparison of different MSD approaches (top) on the same predicted trajectories. The predicted clusters (bottom) of trajectories with FreeTrace, simulated with (*K, H*) ∈ {(0.4, 0.1), (1.5, 0.5), (1.5, 0.99)}, respectively. The empirical MSD (black), variance estimated MSD (blue) for each time lag with the Gaussian fitting, *H* estimated MSD with the Cauchy fitting (red), and the theoretical evolution (green) of molecular diffusion. The empirical MSD (black) corresponds to the ensemble-average of time-averaged square displacements (equation 6) of predicted trajectories. The variance estimated MSD (blue) is the curve where the variance for each time lag is estimated with Gaussian fitting. This approach can provide reliable results only when the number of displacements for each time lag is sufficient, and it becomes highly unstable as the number of displacements reduces as the time lag increases. Our proposed Cauchy fitting (red) estimates *H* without using the MSD-based approach. Since the *H* is accurately estimated for short trajectories at the ensemble level, the expected evolution of molecular diffusion is closer to the theoretical evolution. Given the variance of displacements or square displacement for the first time lag, the evolution of molecular diffusion can be predicted accurately with the estimated *H* with the proposed equation 3. The estimated (*K, H*) values for each population are shown (**c**,**d**,**e**, bottom) with corresponding dots, the theoretical values (green), the estimation with the proposed method (red) and the estimated peaks of the clusters (pink). The total number of trajectories for **c**,**d**, and **e** is 2,300, 2,344 and 2,328, respectively. The mean lengths of trajectories for **c**,**d**, and **e** are 6.61, 6.05 and 5.74 frames, respectively. The error bars correspond to one standard deviation of empirical MSD. The scale bar corresponds to 1 *µ*m with a pixel length of 160 nm.

We present the results of our proposed method (equation 3, 4), the Cauchy fitting for each population, to overcome the bias in *H* estimation at the individual level and the improved accuracy for subdiffusive, Brownian and superdiffusive molecules (**Fig. 3c,d,e**). The proposed method can replace the MSD-based approach. The MSD curves (**Fig. 3c,d,e**, top) are drawn from each corresponding population of clusters (**Fig. 3c,d,e**, bottom). The trajectories are predicted using FreeTrace. The empirical MSD (black, equation 6) is the commonly used approach to calculate the classical diffusion coefficient in the field of SMT. This equation can be highly inaccurate for a small number of non-Brownian particles, and any noise terms are not included. The variance estimation with zero mean Gaussian fitting can also provide MSD. However, this MSD also becomes highly unstable as the number of displacements with a high time lag decreases, and the variance fitting fails severely. Moreover, the variance-based approach for superdiffusive molecules can only be used when the number of trajectories is very high, when their displacement at the ensemble level shows a Gaussian shape. The most critical point of every MSD-based approach is fixing the time point to measure the curvature of MSD represented by the Hurst exponent (*i*.*e*., anomalous diffusion exponent divided by the factor of two). As the number of observed displacements decreases with increasing time lag, fluctuations in MSD make it challenging to choose a time point from a limited number of molecular trajectories in practice.

Our proposed method overcomes the aforementioned difficulties stemming from the MSD-based approach by estimating *H* accurately with a ratio (equation 3). The ratio of displacements eliminates information about the diffusion coefficient, assuming the observed trajectory shows a single regime, and retains only spatio-temporally correlated information about 1D displacement in the ratio distribution. This is particularly beneficial for short trajectories, since motion changes (change of diffusive regime within a trajectory) are less frequently observable. As a consequence, only the estimated variance or square displacement of the first time lag (*i*.*e*., instantaneous diffusion coefficient) and the estimated *H* are utilised to approximate the evolution of MSD, and no fixation of time point is required in our approach. We can observe that the proposed method (red) gives high accordance to the theoretical (green) curve of MSD in **Fig. 3c,d,e** across all types of molecular motion, proving that the proposed method provides more accurate estimates than the MSD-based approach.

We demonstrated that the Cauchy fitting from the predicted trajectories with FreeTrace achieves high estimation accuracy at the ensemble level. The additional validation with simulated trajectories is shown in **Extended Data Fig. 1**. This accuracy is unattainable by the clustering method’s centre finding in short individual trajectories, due to bias from individual estimation. As a consequence, the clusters constructed with FreeTrace can be utilised to classify the individual trajectories as they reveal the trends of *H* and *K*, although with bias. The subsequent estimation of *H* with the Cauchy fitting can provide an accurate approximation of *H* for a single population, even for short trajectories. This signifies that we can accurately measure the temporal correlation of molecular displacements, even for 2-length trajectories (1 ratio for each trajectory), provided the trajectories belong to the same population with equal frame rate for the displacements, and given a sufficient total number of trajectories regardless of length. The total number of trajectories and their average length of each **Fig. 3c,d,e** are 2,300 trajectories with 6.63 frames, 2,344 with 5.43 frames, and 2,328 with 5.49 frames. Note that the estimation accuracy with the Cauchy fitting is not affected by the length of trajectories, but rather the number of observed ratios. The trajectories are simulated without motion changes. Therefore, prior classification of motion-changing trajectories and segmentation [37] are required for experimental data, particularly for long trajectories, before performing the proposed Cauchy fitting.

## Applications to Experimental Datasets

We applied FreeTrace and the proposed Cauchy fitting to three experimental SMT datasets. To test a wide range of conditions, we selected experimental data acquired across different organisms or cell lines, with molecules exhibiting various diffusion properties, using different tagging methods. We selected SMT acquisitions within the cell nucleus as a benchmark to evaluate FreeTrace, as imaging individual molecules in the nucleus is a well-known optical challenge for SMT.

### Chromatin-bound histones (H2B, human cells)

First, as an example of slowly moving molecules, we used previously published SMT data on histone H2B in human osteosarcoma (U2OS) cells [12]. Histones H2A, H2B, H3, and H4 are highly conserved proteins found in eukaryotic cell nuclei. They have two essential functions: packaging the genome and regulating gene accessibility. Since DNA is wrapped around an octamer composed of histones H2A, H2B, H3, and H4, forming chromatin, variations in histone mobility can influence DNA accessibility as well as chromatin stiffness and compaction, with significant consequences for essential biological function [38, 39]. Previous studies revealed the presence of two populations of histones in human cells: histones with low diffusivity (*D <* 0.01*µm*^2^*/sec*), which represent around 75% of the histones and are likely bound to DNA, and histones with high diffusivity (*D >* 2*µm*^2^*/sec*) freely diffusing within the nucleus [40, 41]. Some studies have also revealed the existence of histones switching between immobile and mobile phases [12].

Here, we focus on low-diffusivity histones to gain deeper insight into their diffusive properties. The mobility of H2B was measured by SMT using a U2OS cell line stably expressing H2B molecules fused to Halo, bound to photo-activatable Janelia Fluor 549 dyes (PA-JF549) [42] (**Supplementary movie 1**). The average ratio of the peak of the detected H2B to the background intensity is approximately 3.5. **Fig. 4a** shows trajectories of low-diffusivity histones, predicted with FreeTrace. Using the Cauchy fitting, we were able to recover the Hurst exponent of low-diffusivity histones, reflecting histones incorporated into chromatin. We obtained an estimate of *H* approximately equals to 0.287 (*i*.*e*., *α* ≈ 0.574) from the analysis of all the trajectories from a single cell, consistent with the Rouse model [43] describing chromatin motion, as well as the exponent previously estimated from the MSD of a fluorescent locus inserted into chromatin [44, 12, 45]. This analysis provides the first evidence that the mobility of low-diffusivity H2B histones follows the models described for chromatin in living cells, and the estimated *H* from H2B reveals chromatin’s subdiffusive behaviour indirectly.

**Fig. 4:**
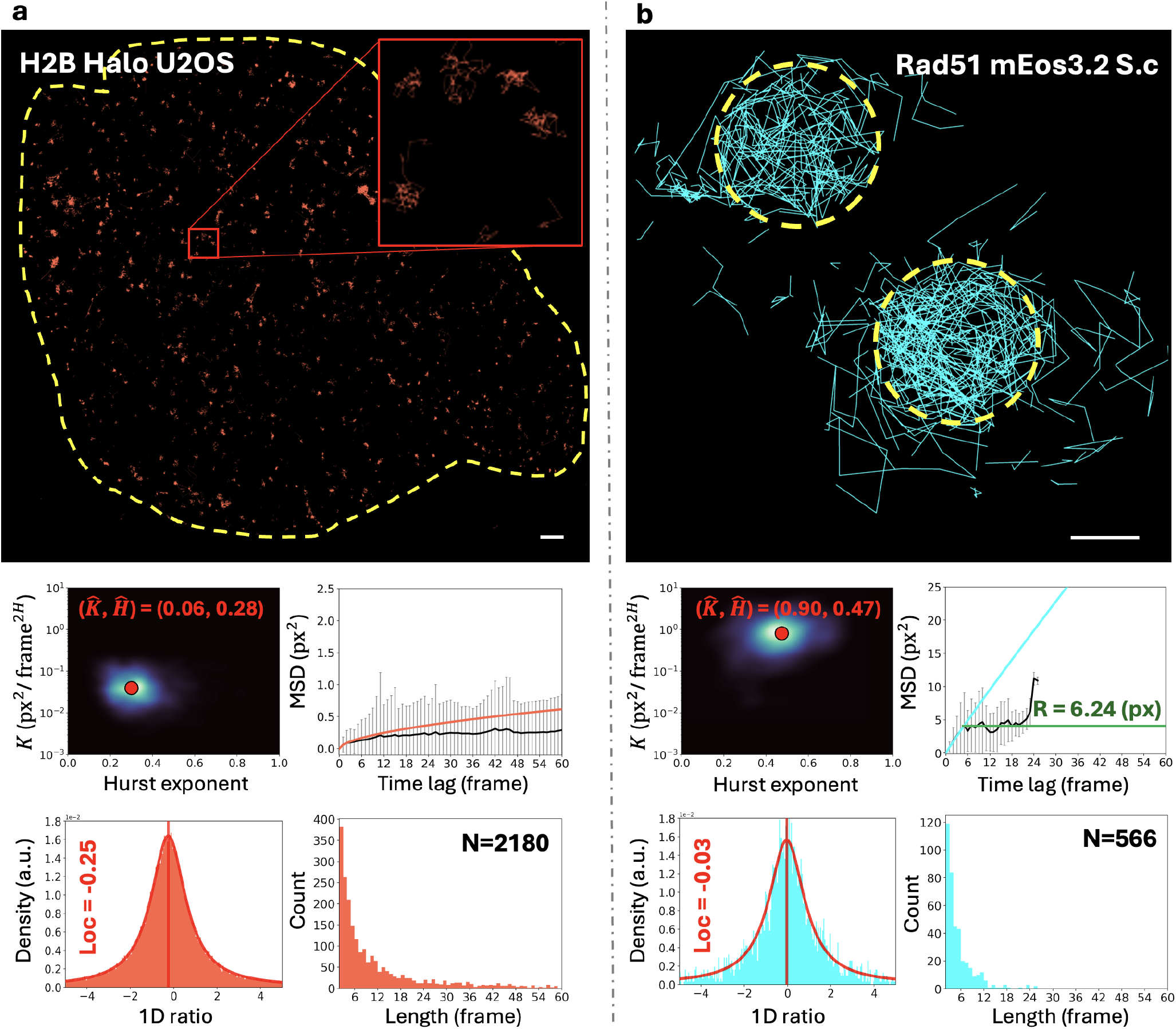
Predicted trajectories with FreeTrace on H2B in human cell and Rad51 in yeast reveal differences between bounded Brownian and subdiffusive motion. **a**, 1st row, The predicted H2B trajectories (red) with FreeTrace in the human nucleus (yellow dashed line). **a**, 2nd row, The cluster of estimated *H* and *K* at the individual level on *H* − *K* space with FreeTrace. The predicted evolution of H2B diffusion is subdiffusive with estimated *H* (0.28) and *K* (0.06) at the ensemble level. **a**, 3rd row, The ratio distribution and the fitted curve (red) with equation 3. The red vertical line indicates the estimated location (−0.25) of the ratio distribution. The estimated location indicates the subdiffusive behaviour of H2B trajectories. The predicted evolution of molecular diffusion (MSD in red) approximately corresponds to 0.021 *µm*^2^*/*sec^0.574^ on the micrometre-second scale. **b**, 1st row, The predicted Rad51 trajectories (cyan) in yeast nucleus. **b**, 2nd row, The predicted evolution of molecular diffusion (cyan) over time in MSD with estimated *H* (0.47) and *K* (0.90) at the ensemble level. The difference between predicted evolution (cyan) and empirical MSD (black) reveals the presence of a nuclear membrane (green) with a radius of approximately 6.24 on pixel scale, corresponding to 0.99 on micrometre scale (bounded Brownian), different from the H2B diffusion (subdiffusive). The radius is approximated by applying the 3-sigma rule. **b**, 3rd row, The estimated location (−0.03) of the ratio distribution indicates that the molecular motion is closer to the Brownian motion. The trajectory-length distribution of trajectories shows the risk of estimating *H* and *K* with the MSD-based approach for a low number of short trajectories. The predicted evolution of molecular diffusion (cyan) approximately corresponds to 0.623 *µm*^2^*/*sec^0.942^ on micrometre-second scale. The trajectories with lengths greater than two frames are included. The total number of trajectories for **a** and **b** is 2,180 and 566, respectively. The empirical means of trajectory lengths for **a** and **b** are 17.17 and 4.86 frames, respectively. The H2B trajectories with estimated *K* values less than 0.2 at the individual level are considered. Pixel size: 160 nm, Framerate: 10 ms (H2B) and 30 ms (Rad51), Video length: 5,000 frames, Scalebar: 1 *µ*m. The error bars correspond to one standard deviation of empirical MSD. The raw images and localised molecules are shown in **Extended Data Fig. 2**.

### Mobility of a repair protein in *Saccharomyces cerevisiae* yeast nucleus

Subsequently, we measured and analysed the mobility of a protein in a microorganism tagged with a different labelling method. We chose to follow the repair protein Rad51 (Radiation sensitive protein 51) endogenously fused to the photo-convertible fluorophore mEOS3.2 in *Saccharomyces cerevisiae* (**Fig. 4b**). The average ratio of the peak of the detected Rad51 to the background intensity is approximately 2.5. We found that Rad51 molecules undergo Brownian motion confined by the nuclear envelope (*H* ≈ 0.471, *i*.*e*., *α* ≈ 0.941). Indeed, the resulting MSD plateau corresponds to a confinement radius of approximately 0.99 *µm*, consistent with the radius of the yeast nucleus. The diffusion motions of H2B and Rad51 illustrate two different types of motions measured in experimental data: subdiffusive motion *versus* bounded Brownian motion. The trajectories in both motions are spatially restricted within a limited time. However, our proposed method distinguishes their motions and provides estimates of *H* even with a small number of short trajectories, across different organismal scales (radii of 1 and 10*µ*m for yeast and human cells, respectively).

### Mobility of a repair protein in human nucleus

Finally, we tested our method on experimental data exhibiting potentially two different diffusion motions. We analysed the mobility of the RNA-binding protein named FUS (human FUsed in Sarcoma). FUS is a multi-functional protein known to regulate transcription, to be recruited at DNA lesions, but also as an essential RNA-binding protein linked to the most severe form of amyotrophic lateral sclerosis (ALS). FUS mobility was quantified using a human cell line expressing FUS endogenously fused to Halo, and bound to photoactivatable dyes (PA-JF549).

The predicted FUS trajectories showed potential clusters (**Fig. 5a**, 3rd row) from the estimated *H* and *K* at the individual level. The first cluster (**Fig. 5b**, cyan) consists of Brownian-like (*H* ≈ 0.52) trajectories with relatively high diffusivity (*K* ≈ 1.08) at the ensemble level. This population is the dominant trajectory type of FUS, accounting for approximately 83% of the sample. The second cluster (**Fig. 5c**, red) showed subdiffusive motion (*H* ≈ 0.17) with relatively low diffusivity (*K* ≈ 0.06). This cluster showed several peaks (**Fig. 5c**, 3rd row), which may indicate that this cluster can also consist of multiple populations. This cluster accounts for 17% of the sample. Note that the motion change is not considered throughout the paper, which causes bias for the estimation of *H* and *K* at the individual level (*e*.*g*., shift of clusters). The boundary (**Fig. 5a**, 3rd row, yellow line) for the clustering is manually chosen based on the following criteria for trajectory classification. **b**: *K* ≥ 0.20, **c**: *K <* 0.20. The average ratio of the peak of the detected FUS to the background intensity is approximately 1.8.

**Fig. 5:**
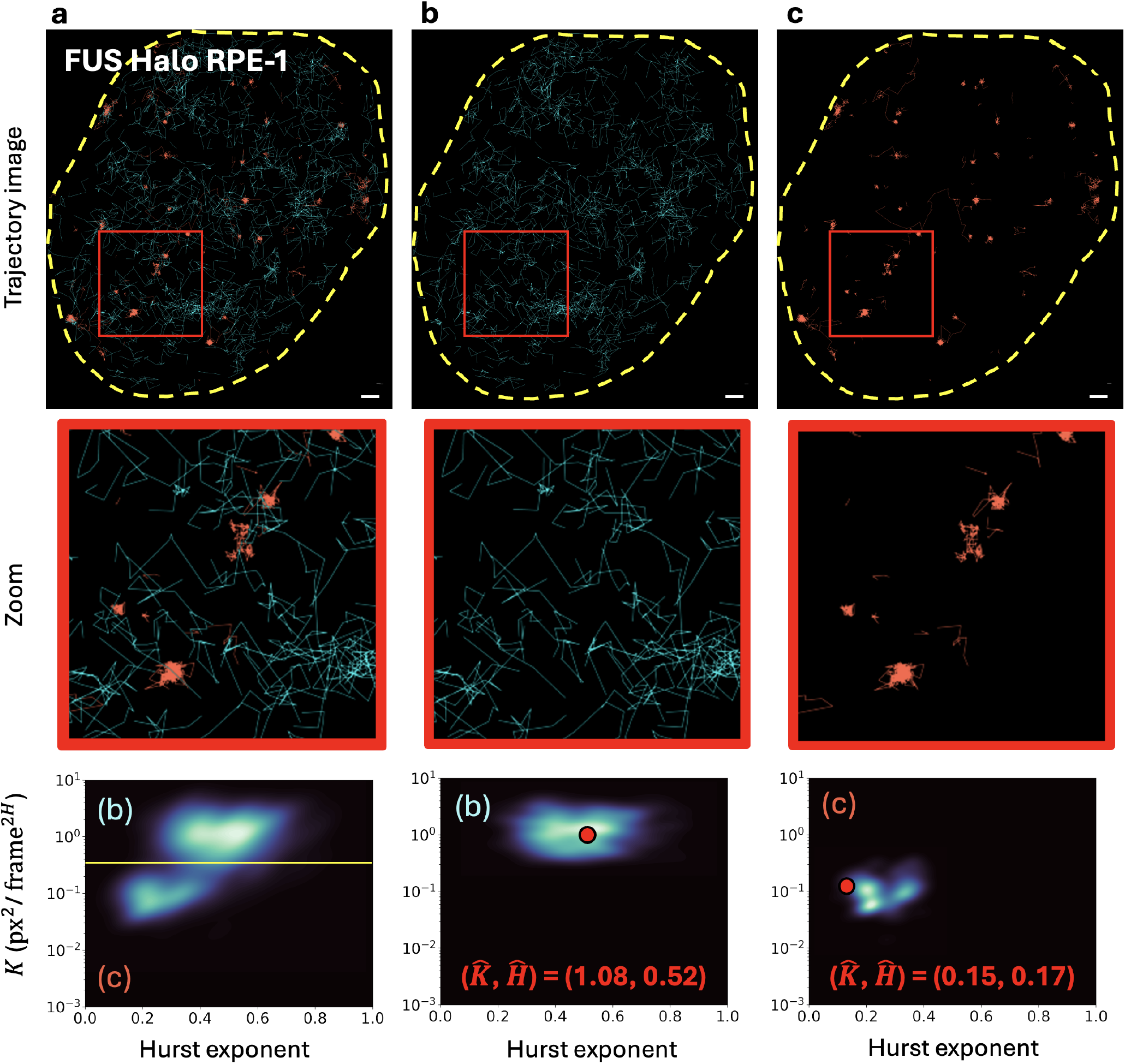
FreeTrace predictions on FUS trajectories in the human nucleus, and the potential clusters of FUS trajectories on the *H*-*K* space. **a**, All FUS trajectories predicted with FreeTrace, with a zoom inside the ROI (red box) and nuclear membrane (dashed yellow line). The predicted trajectories reveal potential clusters (**a**, 3rd row) from the estimated *K* and *H* of individual trajectories. **b, c**, The trajectories classified with respect to the corresponding cluster, divided by the boundaries (**a**, 3rd row, yellow lines). The value of the boundary (yellow lines) dividing the clusters on the *H* − *K* space is as follows: **b**: *K* ≥ 0.20, **c**: *K <* 0.20. **b**, The trajectories (cyan) with higher diffusivity compared to other trajectories, showing a motion closer to Brownian. **c**, The trajectories (red) with low diffusivity and subdiffusive motion. The trajectories with a length greater than two frames are included. The red dots (3rd row) indicate the estimated *H* and *K* at the ensemble level for each cluster with their values. The predicted evolution of molecular diffusion on micrometre-second scale for **b** and **c** is approximately 3.32 *µm*^2^*/*sec^1.04^ and 0.018 *µm*^2^*/*sec^0.34^, respectively. Note that motion-change is not considered, and it induces a bias in *H* and *K* estimation as a result. The distributions and predicted evolution of molecular diffusion in MSD are presented in **Supplementary Fig. 5**. The number of trajectories for each cluster: **a**:1,350, **b**:1,125 and **c**:225. Image size: 134 x 120 px^2^, Pixel size: 160 nm, Framerate: 10 ms, Video length: 5,000 frames, Scalebar: 1 *µ*m.

FreeTrace demonstrated its applicability on real data, with the potential for trajectory classification based on the estimated *H* and *K* values of individual trajectories. The snapshot of raw video, localised molecules, and predicted trajectories are available in **Supplementary Fig. 3b**. The same parameters of FreeTrace are used across all experiments without specifying the maximum jump-distance (**Supplementary Table 1**).

## Discussion

FreeTrace demonstrated the potential for individual trajectory prediction and further classification using the estimated *H* and *K* values of individual trajectories. The improved generality of fBm compared to Brownian motion enables us to analyse the trajectories without modifying any input parameters of FreeTrace, avoiding human bias. FreeTrace proved its performance in the international AnDi challenge [33] as a tool for predicting molecular trajectories from videos. The AI-combined trajectory predictor enabled the estimation of diffusion properties for short trajectories. However, the inevitable bias of the estimated properties for short individual trajectories should be addressed carefully. The bias correction for each individual short trajectory is challenging due to the low number of observations, which gives the future roadmap of trajectory prediction and motion change detection at the individual level. This improvement will allow us to decipher the biological phenomena even with very short and low numbers of trajectories in SMT.

The Cauchy fitting enables the calculation of the spatio-temporally correlated molecular diffusion at the ensemble level under fBm, represented by the Hurst exponent. However, our proposed method (equations 3, 4) does not include any type of noise terms. The method can be further extended by considering various noises, such as localisation noises or noises occurring at the trajectory reconstruction step. We used a non-linear least-squares fitting with L-BFGS-B [46] to estimate the Hurst exponent. We proved that the combination of very accurate estimation of the variance of displacement or the expectation of squared displacement for the first time lag, and the proposed Cauchy fitting-based method can predict the evolution of molecular diffusion at the ensemble level with minimal bias, eventually replacing the MSD-based method. The proposed method requires a sufficient number of ratios regardless of the trajectory length, which can be beneficial in practice, especially for estimating diffusion properties of a set of high-diffusivity molecules. Our proposed fitting function can be applied to the mixture (**Supplementary Fig. 3c**) with further investigation of the optimisation of the loss function.

The estimates of the Hurst exponent revealed the correlated molecular diffusion over time in our experimental data. Interestingly, the estimated *H* values for H2B and FUS trajectories were different: 0.28 and 0.17, respectively. A more accurate *H* estimate for DNA incorporating FUS trajectories can be obtained by segmenting trajectories for motion changes with more trajectories and by experimenting on more cells. However, the difference in *H* may indicate that FUS preferentially interacts with stable regions of the chromatin fibre rather than with homogeneous interactions along the fibre.

The motion-changing trajectories are not considered throughout the paper, even though FreeTrace can produce motion-changing trajectories. However, motion-changing trajectories can be falsely detected, especially in practice with experimental data, due to the current limitation of temporal resolution in SMT. The motion-changing trajectories predicted by FreeTrace can be two independent trajectories whose linkage is mis-reconnected due to the high variance in displacement at the current temporal resolution (*e*.*g*., 10 ms). This uncertainty increases proportionally for high-diffusivity molecules. To identify the time point of motion change, a low molecular density is thus required to reduce false identifications under the current temporal resolution in practice.

The significant increase in temporal resolution at microscopy video acquisition without loss of spatial resolution will be an important breakthrough in SMT. This improvement will yield longer trajectories, enabling precise identification of molecular motion changes and reducing the randomness of molecular diffusion. The reduced randomness naturally leads to a more precise estimation of diffusion properties, and the required number of trajectories for statistical conclusions will be reduced as well. As a consequence, various cell-by-cell biophysical phenomena will be more clearly revealed in live-cell imaging and in SMT, allowing us a better understanding of nature.

## Methods

We utilised default values if any parameter of the algorithms used throughout the paper was not specified. The default input parameters of FreeTrace are utilised throughout the paper except for Δ*T*, which is set to two in **Fig. 2** for objective comparison between software (**Supplementary Table 1**).

The detailed algorithms of FreeTrace, along with the mathematical derivations, are available in the supplementary note. FreeTrace is written in Python, partially with C, utilising Cython. Free-Trace utilises a GPU (NVIDIA) for parallel computations if the system is available. The source code is publicly available in https://github.com/JunwooParkSaribu/FreeTrace. The simulated videos with ground-truth trajectories utilised in this paper are publicly available in https://github.com/JunwooParkSaribu/freetrace_simulation. The versions of the software used in the paper are Track-Mate v7.14.0, TrackIt v1.6, and FreeTrace v1.5.7.

### Human cell culture

#### U2OS H2B-Halo cell line

To visualise the histone H2B, we used a stable U2OS cell line expressing H2B fused to Halo-Tag [12]. Cells were cultured in DMEM (Sigma) supplemented with 10% fetal bovine serum (FBS), 100 *µ*g/ml penicillin, and 100 U/ml streptomycin and maintained at 37°C in a 5% CO2 incubator. For HaloTag labelling, cells were incubated for 30 min with 10 nM of HaloTag ligands conjugated to the photoactivatable dye JF549, kindly provided by the L. Lavis laboratory. Immediately before imaging, the growth medium was replaced with a CO2-independent imaging medium (phenol red-free Leibovitz’s L-15 medium, Gibco). All live-cell experiments were performed on unsynchronised cells.

#### RPE-1 FUS-Halo cell line

To visualise the human protein FUS (FUsed in Sarcoma) by SMT, a RPE-1 cell line was generated by the TACGENE facility (MNHN UMR 7196 - U1154) endogenously expressing Halo-Tagged FUS using CRISPR/Cas9 gene editing system (Kobayashi *et al*., in preparation). All cells were cultured in Dulbecco’s modified eagle medium/nutrient mixture F-12 (Gibco) supplemented with 10% FBS (Gibco) and 100 U/mL Penicillin-Streptomycin (Pen Strp, Gibco) in a humidified incubator at 37°C with 5% CO2. For HaloTag labelling, cells were incubated for 30 min with 200 nM of HaloTag ligands conjugated to the PAJF549, followed by three times wash with PBS (pH 7.4). Samples were kept in imaging medium (phenol red-free Leibovitz’s L-15 medium, Gibco) in the dark till imaging. All human cell lines were tested for mycoplasma using PCR-based assays.

#### Strain and cell culture in *Saccharomyces cerevisiae*

To visualise the repair protein in *Saccharomyces cerevisiae* nucleus, we created a strain expressing endogenous Rad51 fused to mEOS3.2 (plasmid #87030, addgene) in W303 yeast cells following the method described in [47]. Cells were grown at 30°C to early log phase in 4-ml cultures of synthetic culture (SC) medium, and images were taken in a home-made PDMS chamber also maintained at 30°C.

#### Single molecule microscopy acquisitions

All experiments were performed on an inverted Nikon Ti microscope, equipped with an EM-CCD camera (Ixon Ultra 897 Andor) and a 100× 1.4 NA or 1.3 NA oil-immersion objective, leading to a pixel size of 160 nm. Human and yeast cells were maintained at 37°C and 30°C, respectively, using a Tokai device (STXG-TIZWX-SET). For the tracking of single H2B-Halo and FUS-Halo in living human cells, single-molecule imaging sequences of 5,000 frames were acquired at a frame rate of 100 Hz. The PA-JF549 ligand was photo-activated by the 405 nm laser (1 pulsation every 10 frames, power of 0.006 kW/cm^2^ at the sample) and excited by the 561 nm laser (continuous excitation, power of 7 kW/cm^2^ at the sample). For the tracking of Rad51 in the nucleus of living yeast cells, we also acquired sequences of 5,000 frames but at a frame rate of 33.333 Hz, since the transmitted signal of mEOS3.2 is lower than JF549.

#### Simulation of fBm trajectory videos

The motion blur is not taken into account in our simulated videos. However, the motion blur effect becomes stronger for molecules that diffuse in 3D with high diffusivity, and it deforms the PSF and reduces the accuracy of localisations and their precision. As a result, the time gap occurs more frequently in real data than in simulated data without motion blur. This is due to the reduced likelihood of a static Gaussian PSF compared to the deformed PSF of the fluorescence signal in real data. To assess the robustness of the tracking algorithm in each software against the time gap effect causing the unexpected fragmentation of trajectories (*i*.*e*., overestimation of the number of trajectories), we intentionally erased the signals of simulated molecules in the videos. As a consequence, arbitrarily chosen 10% of molecules on average are not visible in the simulated videos used in **Fig. 2**, even though they exist in the ground-truth. Hence, no software can detect 10% of localisations on average across all simulated videos for the test of unexpected trajectory fragmentation. The 10% (proportional to the length) of molecules are chosen uniformly across the videos. Variables of trajectories (**Supplementary note 1**) measured with elapsed time greater than 1 frame are divided into two sub-trajectories. This division is not accounted for in the predicted total number of trajectories. We therefore processed the time gap equally for all predicted trajectories of each software. The average SNR (Signal-to-noise ratio) of the fluorescence signal in the simulated videos is 3. Any additional filtering or post-processing of trajectories is not applied throughout the paper.

#### Detection of moving particles with likelihood ratio and subsequent sub-pixel localisation

FreeTrace consists of three major and one optional algorithm (**Fig. 1c, Supplementary Note 2**). First, it performs single particle (molecule) detection in a given video using a likelihood ratio test [3]. We assume the background is contaminated with Gaussian noise (approximately Poisson noise of high photon counts). The likelihood ℒ_1_ (equation 1), an isotropic 2D Gaussian function with radius *r* inside square sliding windows of length *w*, is maximised with respect to *m*_1_, *σ*_1_ and *I*, where *m*_1_, *σ*_1_ are the mean and SD (standard deviation) of Gaussian noise in the presence of a Gaussian PSF (point spread function) at the centre of sliding window. The likelihood ℒ_0_ is also maximised at the same sliding window with respect to *m*_0_ and *σ*_0_, which correspond to the mean and SD of Gaussian noise without the presence of a Gaussian PSF.

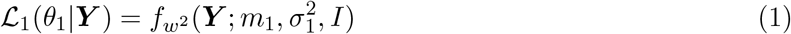

The ratio of ℒ_1_ and ℒ_0_ generates a view of likelihood ratios which acts as a 2D Gaussian PSF filter **Supplementary Fig. 1a**). We collect only the PSF satisfying the *λ* (detection threshold) with AIC. The subsequent 2D Gaussian fitting (equation 2) provides the estimated sub-pixel positions 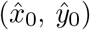, xy variances 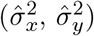, intensity factor (*Î*) and spatial correlation 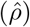 of empirical PSF.

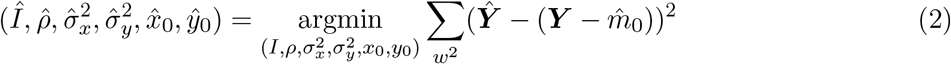

The image inside a sliding window is subtracted beforehand by the estimated mean of the background signal 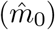 for the simplicity of the computation. Note that determining the precise position of a molecule for a given time interval is nearly impossible since the molecule is moving during the acquisition. Moreover, the deformation of the PSF in SMT is frequently observed, as the molecule continuously moves in 3D space during acquisition. This deformation becomes stronger for fast-diffusive molecules, and we can no longer accurately approximate the position of the molecule as in usual SMLM (Single-molecule localisation microscopy) on nanometre scale, which is a physical limitation of SMT. We therefore did not perform the estimation of localisation precision as in SMLM. Readers should note that the estimated position of the molecule in SMT is an approximation of one of the probable positions the molecule travels along during the acquisition time, which is one of the factors contributing to the displacement variance in the molecular trajectory. The detailed detection and localisation of single molecules from microscopy video is available in **Supplementary Note 3**.

#### Reconnection of detected molecules under fBm

For computational efficiency and simplicity of molecular trajectory reconstruction algorithms, the reconnection of localised molecules in FreeTrace is built on the following assumptions, under which it exhibits optimal performance.

##### Assumption 1

Given a set of molecules, each molecule follows fBm with a fixed *H* (Hurst exponent) and *K* (generalised diffusion coefficient).

##### Assumption 2

The motion of each molecule is isotropic in all dimensions. (*i*.*e*., *H* and *K* is equal in each dimension)

##### Assumption 3

The *H* of the given system is bounded between 0 and 1, exclusive.

The JD_*max*_ is the maximum displacement possible that molecules can move in 2-dimensional space for a given frame rate. FreeTrace estimates JD_*max*_ if it is not given by the users. Since the increment of one-dimensional fBm is a stationary, non-independent Gaussian distribution, we approximate the JD_*max*_ with a Gaussian mixture model. The displacements of the nearest-neighbour between each consecutive frames are sampled to approximate the JD_*max*_. We test different numbers of populations using the Bayesian information criterion (BIC) to determine the optimal number of populations. Note that this process is highly unstable for the low number of trajectories, as the mixture fitting can potentially overfit locally, resulting in high bias. We fixed the minimum JD_*max*_ as 5 pixels (800nm for a 160nm pixel length) to avoid a scenario where a small number of fast diffusive populations is unfitted due to the large number of slow populations. The JD_*max*_ not only determines the maximum distance that a molecule can travel, but it also reduces the computational cost of reconnections by reducing the size of the directed acyclic graph (DAG). This estimation is skipped by manually setting the JD_*max*_.

The important property of fBm is the spatio-temporal correlation of increments (*i*.*e*., displacements) of the trajectory. This property is not easily observable in the short range and in the distribution of displacements. However, since we want to observe the temporal correlation of trajectories and apply the correlation for the reconnection step, we used a ratio distribution of displacements (equation 3) and its quantile function (equation 5) as cost functions. We construct a DAG (**Supplementary Fig. 1c**) of a depth Δ*T* between the localised molecule within the range of JD_*max*_. We did not construct a bipartite graph due to the event where the fast molecule is not detected with low signals, including the blinking event. This event is inevitable because decision-making is binary, and it becomes more frequently observable as the diffusivity of the molecule increases. This problem is addressed using the *gap* strategy in other software. The edges of the graph are weighted with the log density with respect to Δ*t*, Δ*s* and *Ĥ*. The detailed derivations from the fBm trajectory to the Cauchy distribution, along with the algorithms, are available in **Supplementary Note 4**. Note that we estimate *H* and *K* with neural networks with Convolutional LSTM (Long Short-Term Memory)[48] for individual trajectories, which has already proven its effectiveness [37]. The feedback loop continuously estimates *H* of past trajectories over frames to weight the edge of the DAG for the reconnection of future trajectories.

The density function of the increment ratio of the fBm trajectory is as follows:

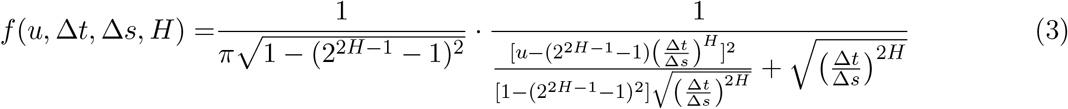

where *u* is the ratio of 1D displacement, Δ*t* and Δ*s* are elapsed time between two displacements and *H* is the Hurst exponent. The values of equation 3 with respect to different *H* are shown in **Supplementary Fig. 1d**.

The distribution function of equation 3 is as follows:

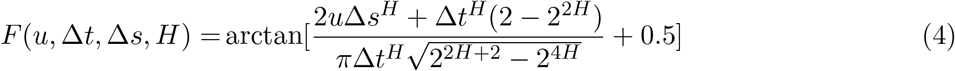

The quantile function of equation 4 is as follows:

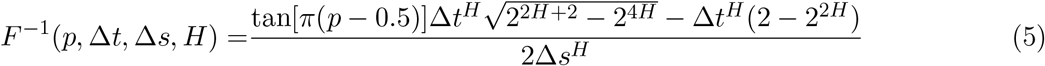

where *p* is the probability. Note that *H* is bounded between 0 and 1, exclusive.

### Cauchy fitting for the estimation of Hurst exponent at the ensemble level

Equation 3 describes how the spatio-temporal correlation of molecular diffusion at the ensemble level is revealed in the ratio distribution with respect to the Hurst exponent. Suppose the ratio of frame rates is the same for a given sequence. In that case, the ratio distribution of Brownian molecules converges to the standard Cauchy distribution with a location at zero. In contrast, the location of the ratio of subdiffusive molecules converges to −0.5 as lim_*H*→0_ (**Supplementary Fig. 1d**). The positive spatio-temporal correlation indicates that the location converges to 1.0 as lim_*H*→1_. From equations 3 and 4, we fit the Cauchy distribution to the empirical ratio distribution to estimate the Hurst exponent of a given set of molecules at the ensemble level. We utilised the non-linear least squares method for the fitting, assuming an equal frame rate for the trajectories.

This fitting is performed on the distribution where the information of the generalised diffusion coefficient is eliminated. The MSD-based [19] approach requires a fixation of a time-point for calculating the *H* and *K* ensembles, which necessarily discards a significant amount of data as the fixed time-point approaches zero. However, this fitting utilises all data as a displacement fitting for estimating the diffusion coefficient or the variance of displacements. As it is performed at the ensemble level, the bias, which comes from the *H* estimation using a clustering-based method on the individual level with a neural network, is significantly lower than that of the proposed methods [33].

Note that we performed the Cauchy fitting on the truncated ratio distribution with a range of −10,000 to 10,000. Prior segmentations and classifications of individual trajectories are required for accurate estimation in cases involving 2 or more populations with motion-changing molecules. The Cauchy fitting can be extended to 2 or 3 populations with further investigation of optimisation to approximate the ratio of populations for a given sample (**Supplementary Fig. 3c**). The Cauchy distributions with respect to *H* values are shown in **Supplementary Fig. 1d** with their derivation in **Supplementary Note 4**.

### Mean squared displacement

The ensemble-averaged and time-averaged squared displacement without noise for the elapsed time (time lag) *τ* for a given set of 1D trajectories is as follows:

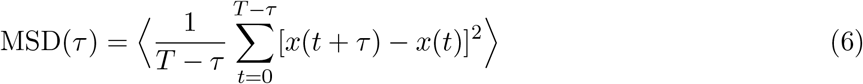

where ⟨⟩ denotes the ensemble-average, and *x*(*t*) corresponds to the x coordinate of a molecule at time *t*, which follows a power law function for fBm *asymptotically* as described below,

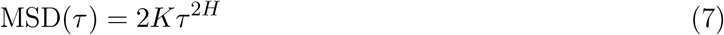

where *K* is the generalised diffusion coefficient and *H* is the Hurst exponent.

### Estimation of *H* and *K* at the individual level

We employed the architecture of neural networks presented in [37] for estimating *H* and *K* for each individual trajectory. The network for *H* estimation is trained with Convolutional Long Short-Term Memory (ConvLSTM) [48], and the network for *K* estimation is trained with a relatively simple neural network trained on the log space with displacements of trajectories. The detailed architecture and inputs of neural networks are available in the aforementioned paper.

## Acknowledgements

We thank C. Chaumeton, and F. Lam from the Cellular Imaging Facility (Institut de Biologie Paris-Seine, Sorbonne Université) for their support on the SMT microscopes, as well as the ARTBio facility for providing data storage. We also would like to thank A. Brion and J-P Concordet from the TACGENE facility (U1154–UMR7196) for the creation of the RPE-1 cell lines expressing endogenous FUS-Halo. JMH team received financial support from the Agence Nationale de la Recherche (ANR-22-CE12-0039 AROSE), the i-Bio Initiative from the Idex Sorbonne University Alliance, the IBPS Incentive Action and the ATIP Avenir 2021.

## Declarations

The authors declare no competing interests.

## Figure

**Extended Data Fig. 1:**
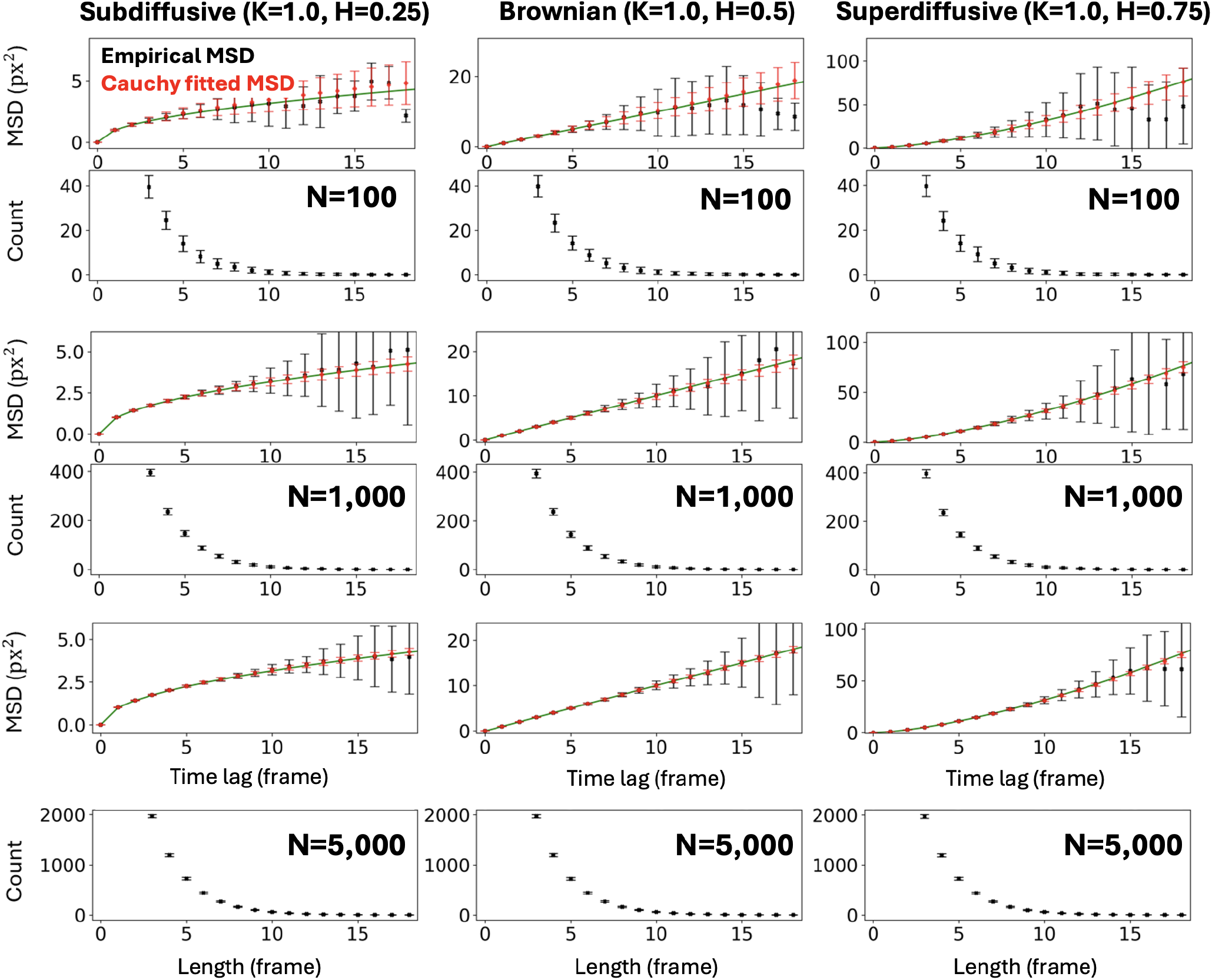
The Cauchy fitting for predicting the evolution of molecular diffusion across 100 repeated fBm simulations. The comparison between the Cauchy fitting and empirical MSD on 100 simulations, varying the number of short trajectories. Each MSD is shown with the corresponding length distribution of trajectories. The trajectories with lengths greater than two are included in the MSD. Only the localisation noise is accounted for in the figure, and the mis-reconstruction noise is not included. The coordinates of trajectories are contaminated with zero-mean Gaussian noise of standard deviation 0.1 pixel. The Cauchy fitting proves effective for estimating *H* for short trajectories across all scenarios. The estimated temporal evolution of molecular diffusion can be approximated with the Cauchy fitting, particularly effective for short trajectories. The scenarios, from left to right, correspond to subdiffusive, Brownian, and superdiffusive motions. The number of molecules for each scenario is 100, 1,000 and 5,000 from top to bottom. Each scenario is repeated 100 times. The error bars indicate the 1 standard deviation of MSD from 100 simulations.

**Extended Data Fig. 2:**
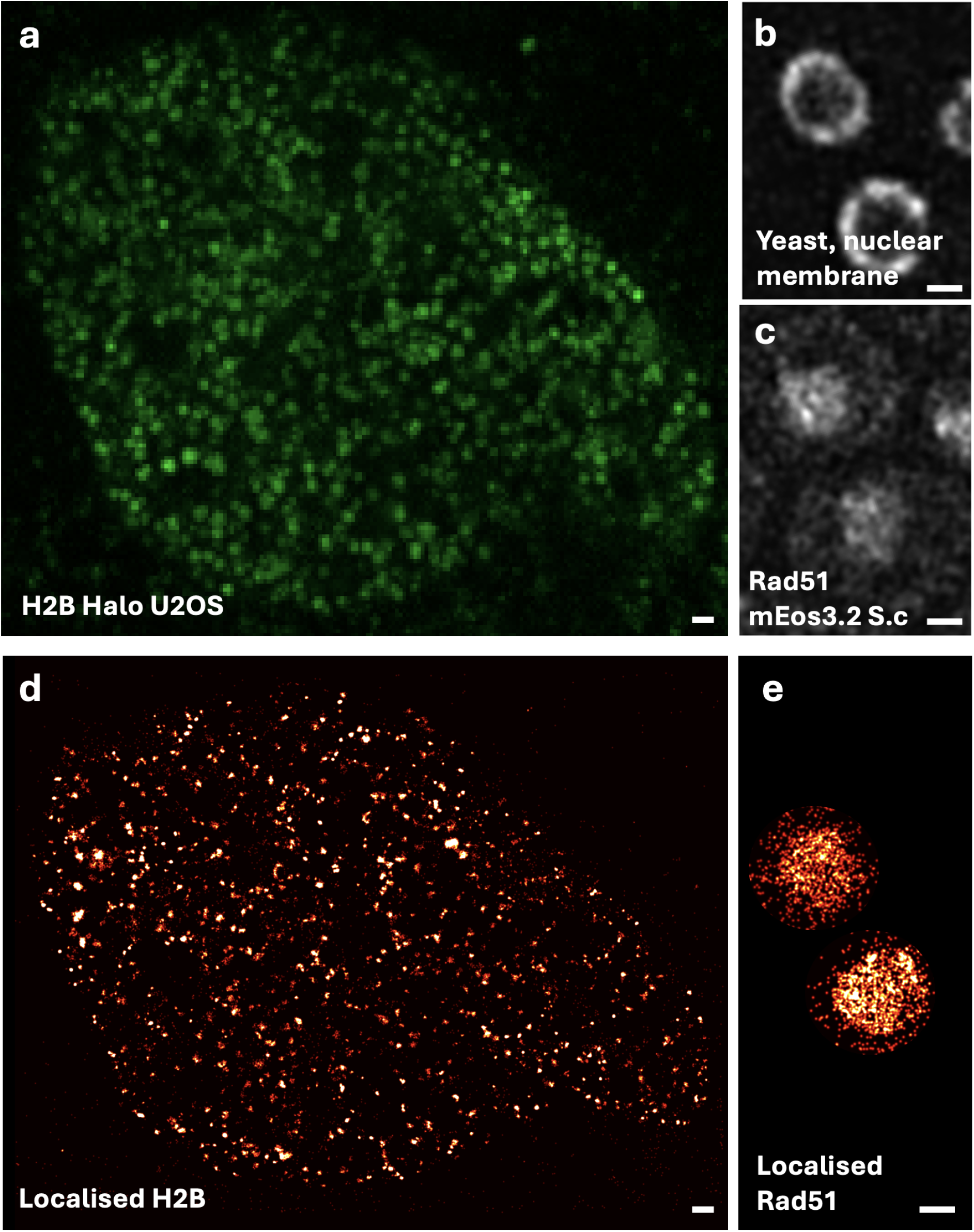
The raw and localised H2B molecules in a human cell and Rad51 molecules in budding yeast cells with FreeTrace. **a**, The Z-projected maximum intensities of H2B signals in a human cell. **b**, The nuclear membranes of yeast cells. **c**, The Z-projected maximum intensities of Rad51 signals in yeast cells. **d**, The localised single H2B molecules in **a** with FreeTrace. **e**, The localised single Rad51 molecules in **c** with FreeTrace. The reconstructed trajectories and their diffusion properties are presented in **Fig. 4**. The localised molecular density is presented in colours ranging from black to white in **d** and **e**. The scale bar indicates 1*µm*. The video length is 5,000 frames for both. The size of ROI in yeast is slightly larger than its nuclear membrane, and the ratio of the number of reflection events at the nuclear membrane to the number of free-diffusive Rad51 in the nucleus affects the estimate of *H*.

## Supplementary information

**Supplementary Fig. 1:**
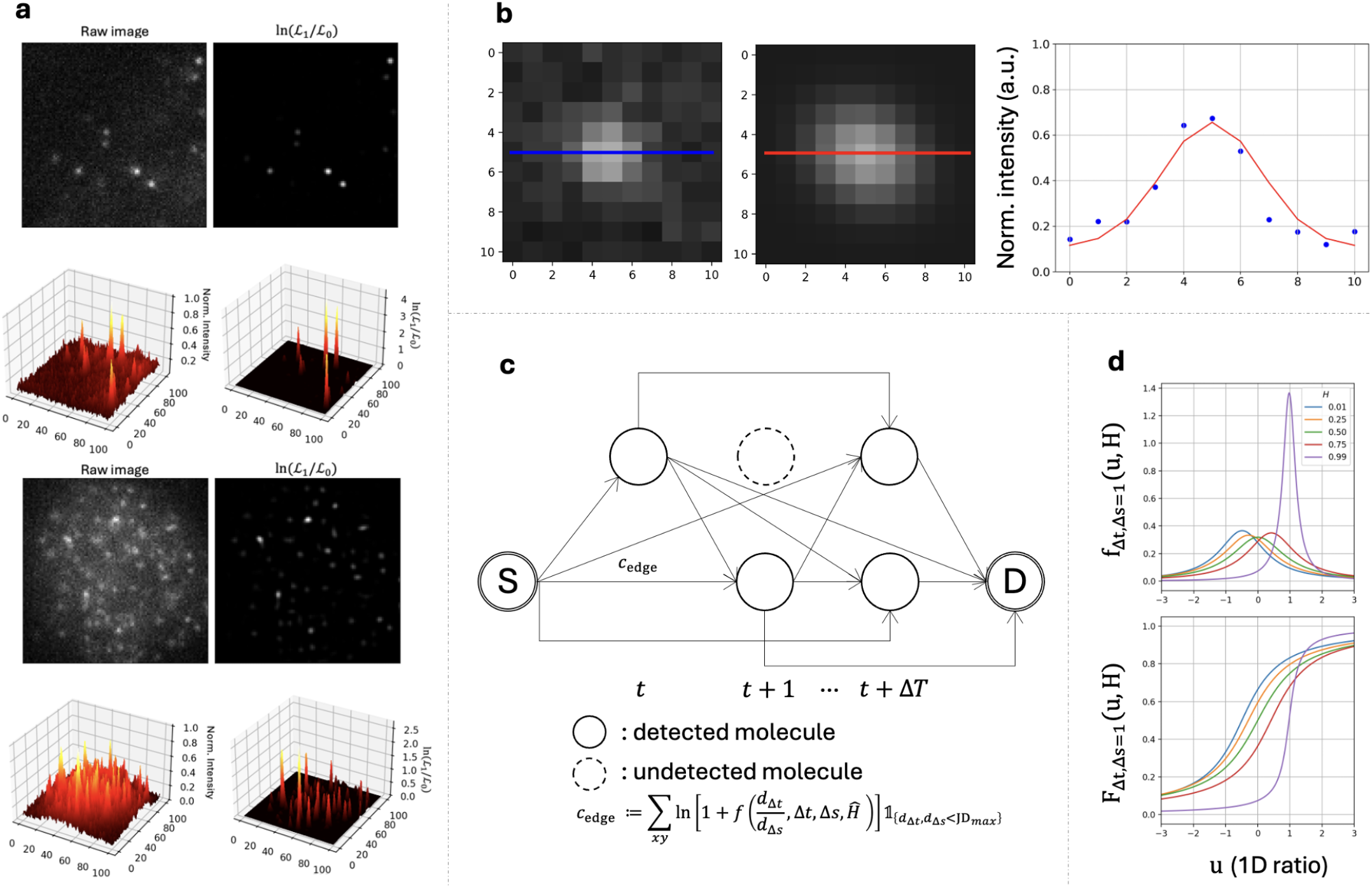
The likelihood ratio test and regression to localise fluorescence signal, the graph for the reconnection of detected molecules, and the graphs of the Cauchy distributions with respect to *H*. **a**, The high and low-SNR images were mapped with the likelihood ratio, respectively. The left images show the raw images in 2D (top left) and 3D (bottom left), while the right images show the log likelihood ratio between the fluorescence signal and background in 2D (top right) and 3D (bottom right). The likelihood ratio of the fluorescent signal in low SNR is lower than that of high SNR due to the heterogeneous background signal. Equation 1 gives the likelihood of a fluorescent signal, and the comparison over background likelihood gives a filtered image as shown in the right side images. **b**, Fitted 2D Gaussian function (mid) (equation 2) on empirical observations (left) inside a square window of length 11. The curve of the figure (right) shows the values at the centre horizontal line for the observations (blue) and estimations (red), respectively. The sub-pixel values (*x* and *y* coordinates of the peak of the fitted 2D Gaussian function) correspond to the 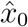 and *ŷ*_0_ in equation 2. **c**, Schematic representation of a local DAG (Directed Acyclic Graph) from the source node to the destination node. Both source and destination nodes are dummy nodes used to link intermediate observations, ensuring the completeness of the graph. The intermediate nodes are the molecules from *t* to *t* + Δ*T* with edges weighted with *c*_edge_, which is a density function (equation 3) of a Cauchy distribution. The estimated *Ĥ* of current trajectories are passed to the next graph for the computation of the next *c*_edge_. **d**, Theoretical Cauchy density (upper) and distribution (lower) function of fBm trajectories with respect to the displacement ratio *u* and Hurst exponent *H* when time intervals Δ*t* and Δ*s* are equal for each consecutive displacement.

**Supplementary Fig. 2:**
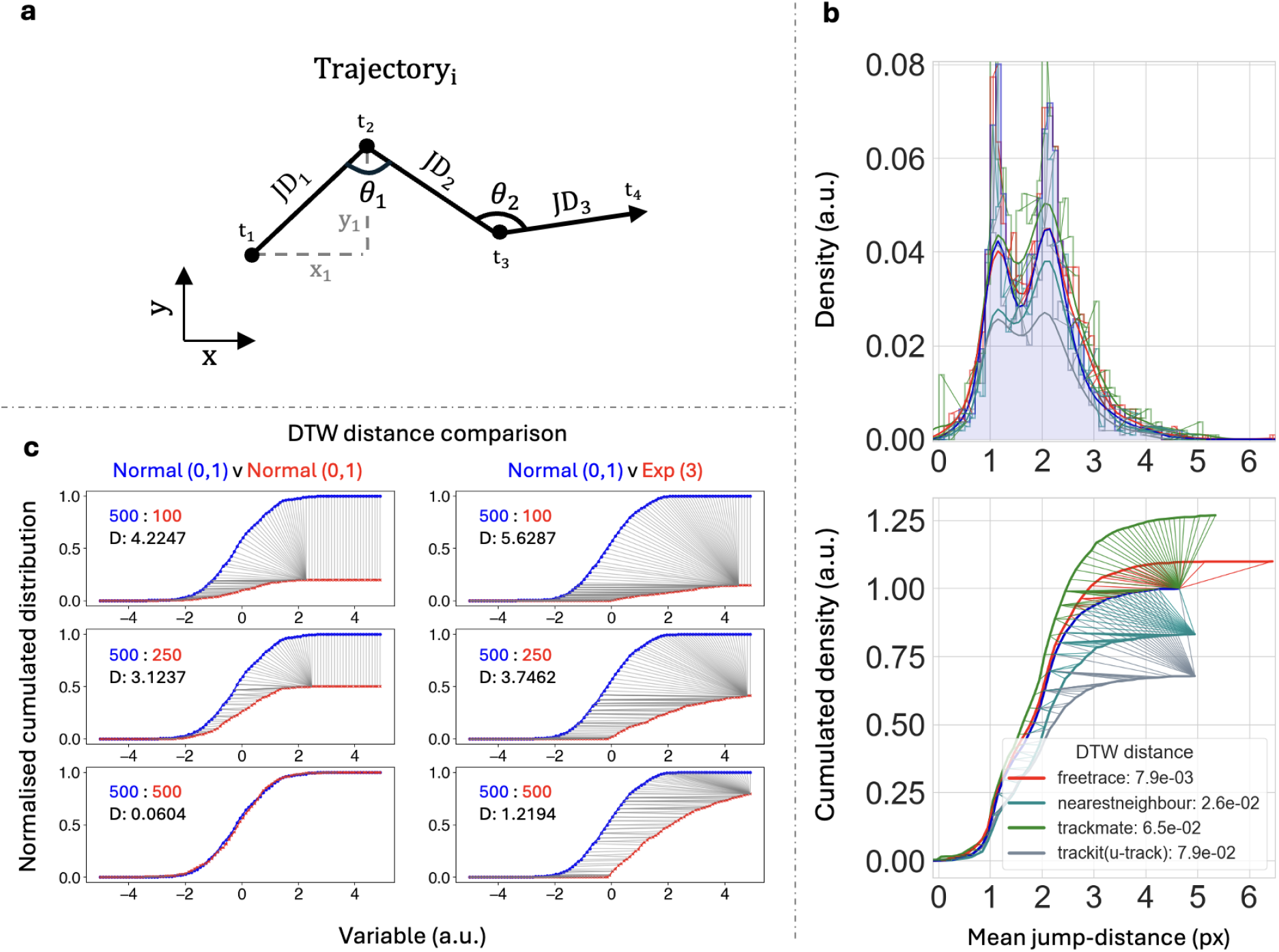
The overview of DTW distance and variables used in the paper with graphical representation. **a**, Presentation of the variable (*X*_*i*_) which can be obtained from 2D trajectory. 2D displacements are a set of all JD (jump-distance) in the entire sample. Mean jump distances are a set of the averaged JD for each individual trajectory; hence, the cardinality is equal to the number of predicted trajectories. 1D displacements are a union of two sets, x and y displacements (they will be the same as the number of trajectories if the sample size is infinite, assuming isotropic molecular motion). **b**, The result of mean jump distances at low density (**Fig. 2**) between software with the best paths (coloured bridges between curves) obtained with DTW on the PDF (Probability Density Function, top) and CDF (Cumulative Distribution Function, bottom). The distances in legend are multiplied by the distance from the density function and from the cumulative distribution. The blue curve corresponds to the GT of the scenario shown in **Fig. 2** at low molecular density. The PDF and CDF of each software are normalised to the GT. **c**, The result of DTW distances compared between two normal distributions or between the normal distribution and the exponential distribution with respect to different numbers of observations. As the number of observations (red) matches the reference number of observations (blue), the DTW distance approaches 0. We used the DTW distance to measure the difference between two observations, taking into account the predicted number of observations for each software.

**Supplementary Fig. 3:**
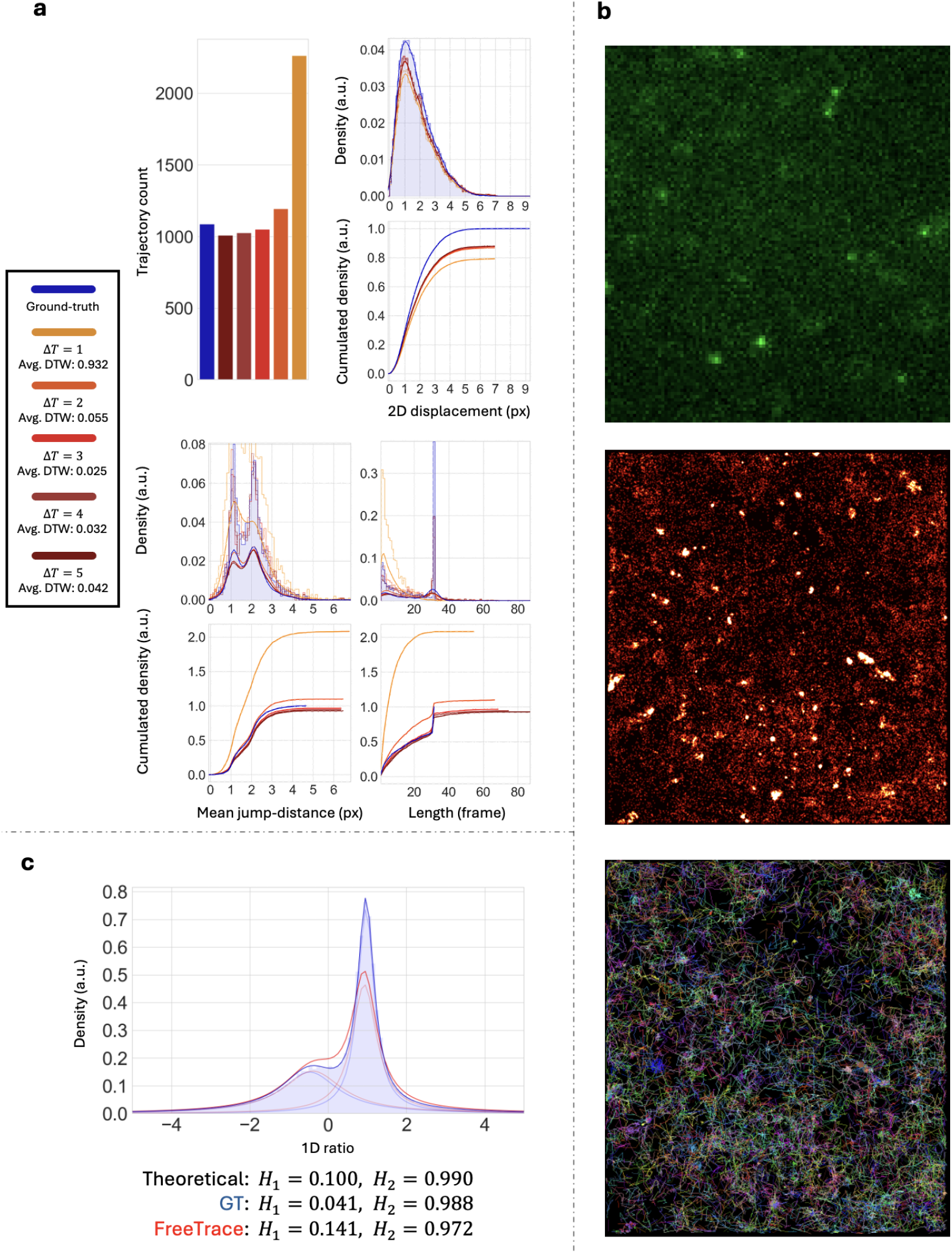
Depth of the graph Δ*T* and ratio fitting for the estimation of the Hurst exponent. **a**, The change of FreeTrace performances with respect to Δ*T*, the depth of graph at the reconstruction step, on the scenario presented in **Fig. 2a**. FreeTrace showed the best result at Δ*T* = 3 at low density. The increased Δ*t* leads to an increased computational time, where the complexity is factorial depending on the molecular density of neighbours (**Supplementary Fig. 1c**). **b**, The snapshot of FUS video, localised molecules, and the predicted trajectories presented in **Fig. 5**. The density map of localised molecules (**b**, mid), ranging from black (low density) to white (high density). The colours (**b**, bottom) indicate the individual predicted trajectories. The localised and the predicted trajectory images are given as the default outputs of FreeTrace. **c**, The predicted ratio distribution and its Cauchy fitting on GT (blue) and FreeTrace (red) on the simulated 2-population sample (1,644 trajectories) with *H* = 0.1 and *H* = 0.99. FreeTrace accurately estimates the number of subdiffusive ratios. However, it underestimates the number of superdiffusive ratios. The mixture of equation 4 is utilised with non-linear least square fitting. Since the ratio distribution does not include the information of (generalised) diffusion coefficient, it only reveals the correlation of molecular movements.

**Supplementary Fig. 4:**
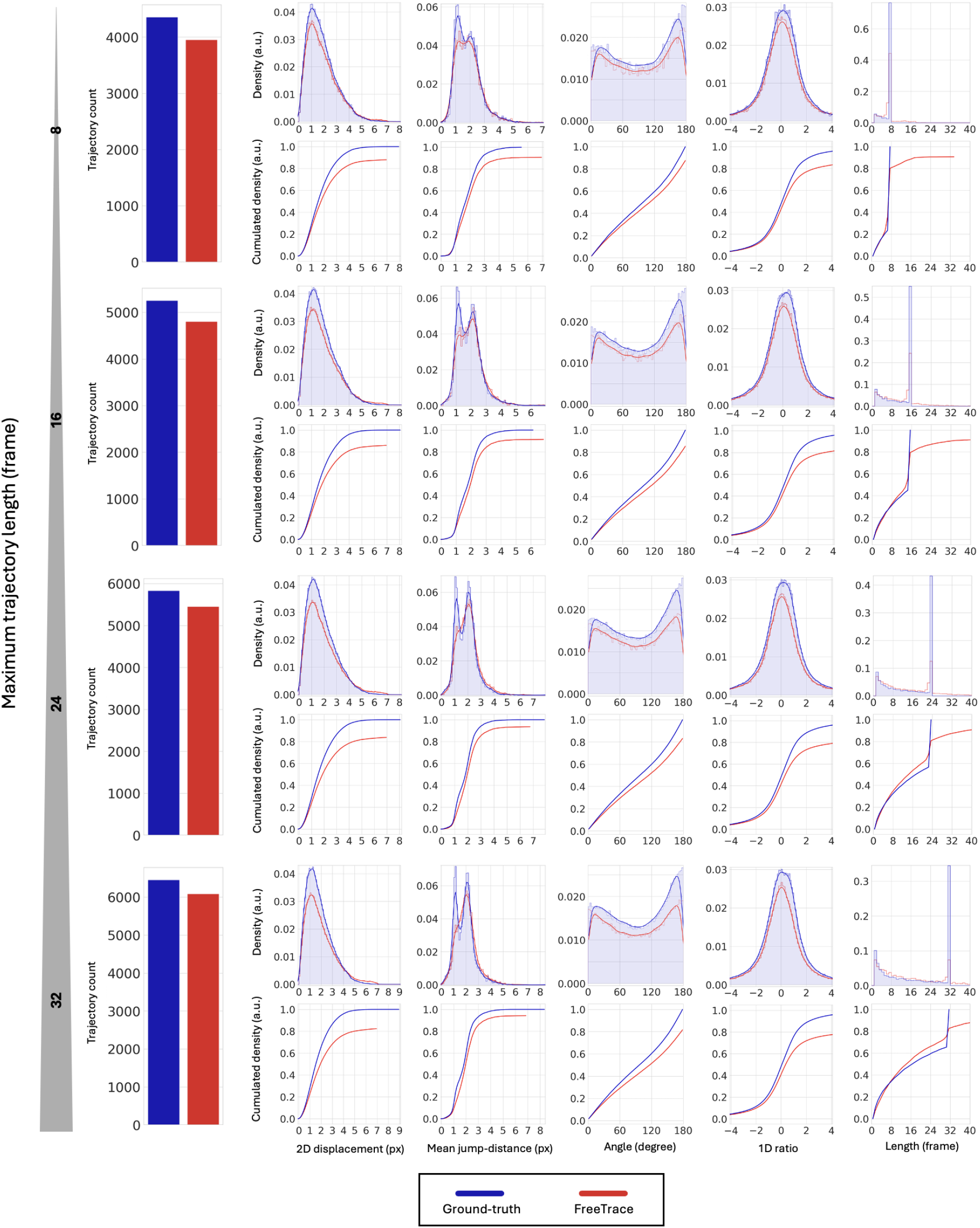
Performance of FreeTrace depending on the length of trajectories. The performance of FreeTrace is predicted on the simulated videos with respect to different lengths of molecular trajectories. The same scenario of *K* and *H* is used as shown in **Fig. 2** at high density. The Δ*T* = 3 is used for the construction of the reconnection graph in the prediction compared to **Fig. 2**. Note that the molecular density increases as the length of the trajectory becomes longer, showing the high density at 32 length, and 10% of molecules are dropped as the simulated scenario in **Fig. 2**. The mis-reconnection is observable and increased, as the length of the trajectory becomes longer, in the mean jump-distance histogram. This is natural since the probability of mis-reconnection increases as the trajectory remains within the field of view (FOV) for a longer period.

**Supplementary Fig. 5:**
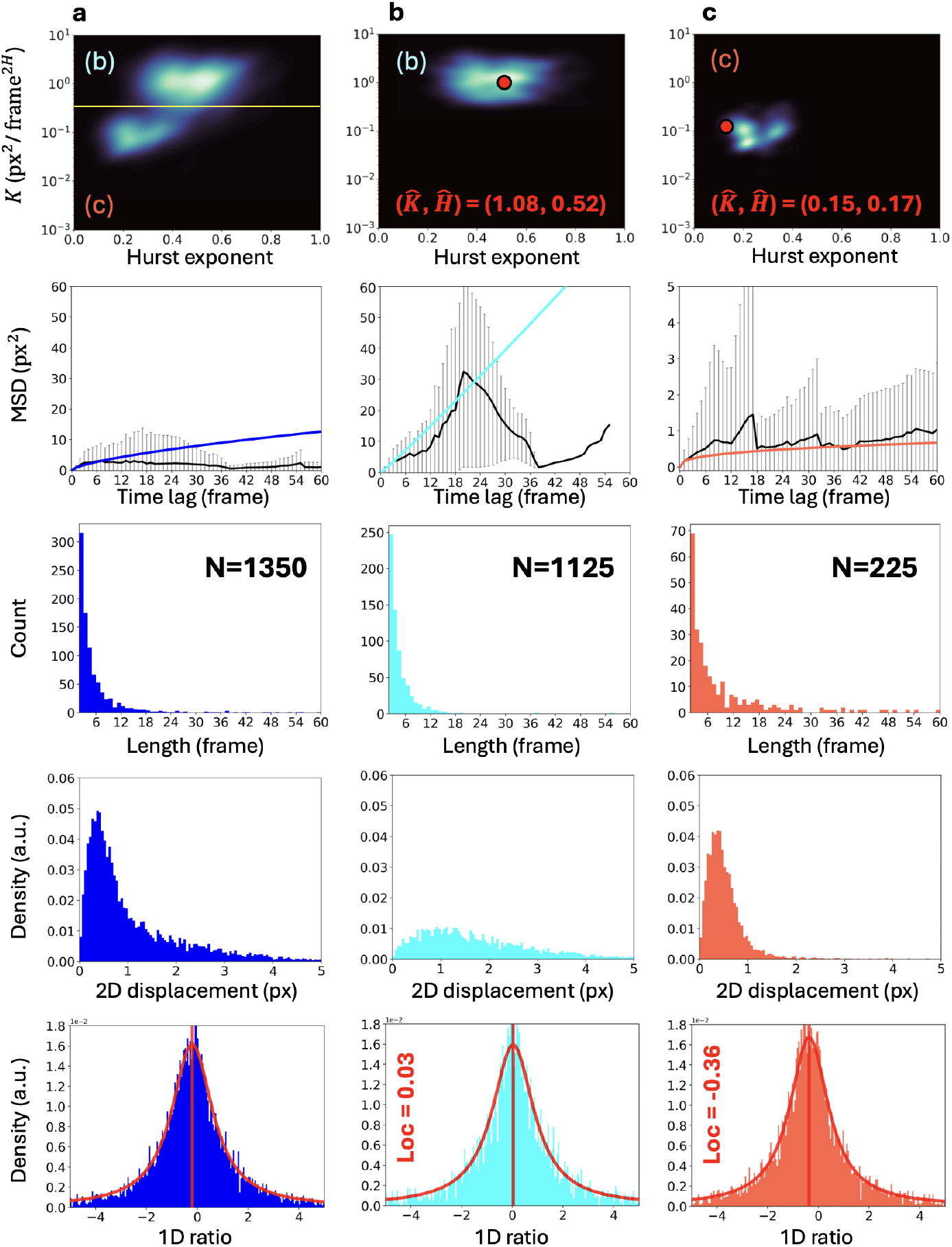
The distributions of the predicted FUS trajectories in each cluster. The clusters (1st row) of estimated *K* and *H* of all individual trajectories (**a**) and the clusters of the trajectories in the divided regions **b** (cyan), **c** (red) as shown in **Fig. 5**. The red dots indicate the estimated *K* and *H* values for each cluster at the ensemble level. The empirical MSD (2nd row, black) and the evolution of MSD using the proposed method for each cluster are shown along with the estimated *K* and *H* for each cluster. The distribution of trajectory length (3rd row) in the corresponding cluster. The 2D displacement distributions (4th row) of the trajectories in the corresponding cluster. The 1D ratio distributions (5th row) of the trajectories and fitted curves with estimated locations. **a**, The distributions of FUS trajectories before classification with the boundary (yellow line) on *H* − *K* space. **b**, The trajectories with the estimated *H* value closer to 0.5 at the ensemble level, indicating a motion closer to Brownian with fast diffusivity compared to the trajectories in **c**, and it is the dominant population in the sample. **c**, The cluster with several peaks at the individual level with the estimated *H* value around 0.17 at the ensemble level, representing the subdiffusive motion with slow diffusivity. The peak of the cluster is possibly biased at the individual level, as shown in **Fig. 3c**, due to the short length of trajectories. This population is the least dominant in the sample. The motion changes in trajectories are ignored when estimating *H* and *K* at the individual level, which causes bias in the individual-level estimations. The small clusters in **c** may be caused by the bias of *H* estimation at the individual level, which needs segmentations of trajectories for a more reliable decision. The estimated value (*K, H*) (red dot) for each cluster: **b**: (1.08, 0.52), **c**: (0.15, 0.17). The trajectories where the length is greater than two frames are included. The number of trajectories for each cluster: **a**:1,350^6^, **b**:1,125, **c**:225. The corresponding trajectories for each cluster are visualised in **Fig. 5**.

**Supplementary Table 1:**
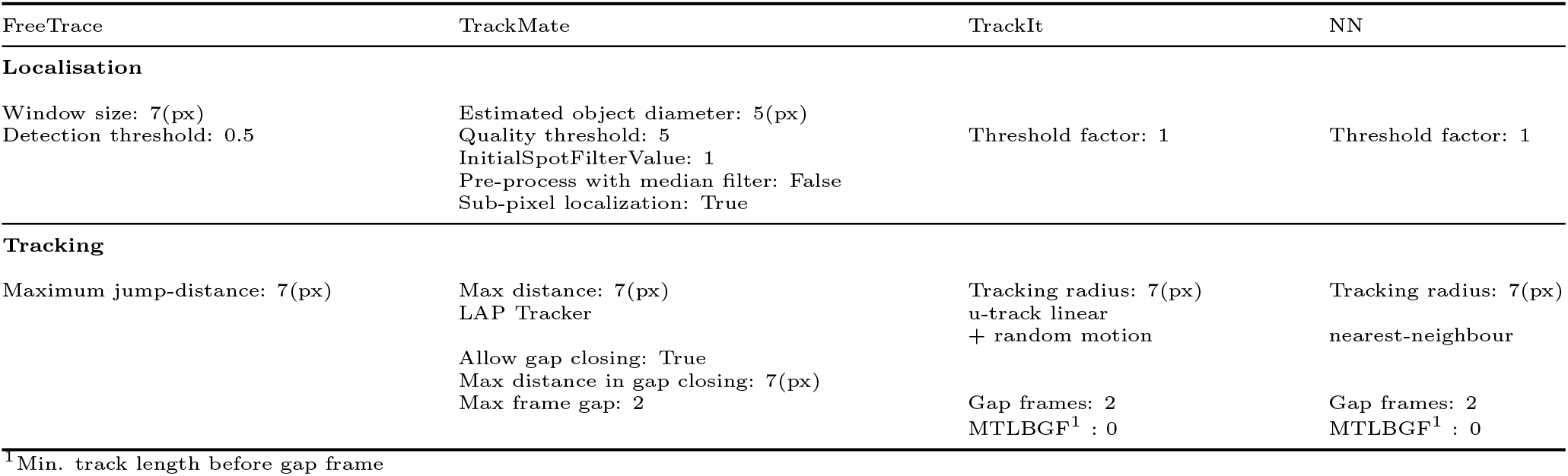
The parameters utilised for each software for the results of **Fig. 2** in the main manuscript. Any parameters or algorithms not specified in this table were set to the default values or not utilised.

| FreeTrace | TrackMate | TrackIt | NN |
| --- | --- | --- | --- |
| **Localisation** |  |  |  |
| Window size: 7(px)<br>Detection threshold: 0.5 | Estimated object diameter: 5(px)<br>Quality threshold: 5<br>InitialSpotFilterValue: 1<br>Pre-process with median filter: False<br>Sub-pixel localization: True | Threshold factor: 1 | Threshold factor: 1 |
| **Tracking** |  |  |  |
| Maximum jump-distance: 7(px) | Max distance: 7(px)<br>LAP Tracker<br>Allow gap closing: True<br>Max distance in gap closing: 7(px)<br>Max frame gap: 2 | Tracking radius: 7(px)<br>u-track linear<br>+ random motion<br>Gap frames: 2<br>MTLBGF<sup>1</sup> : 0 | Tracking radius: 7(px)<br>nearest-neighbour<br>Gap frames: 2<br>MTLBGF<sup>1</sup> : 0 |

<sup>1</sup>Min. track length before gap frame

## 1 Variables of trajectory in SMT

The variables (**Supplementary Fig. 2a**) utilised throughout the paper are as follows, assuming the isotropic trajectories and the variables of trajectories have an equal time lag (*i*.*e*., each variable is measured during the same elapsed time).

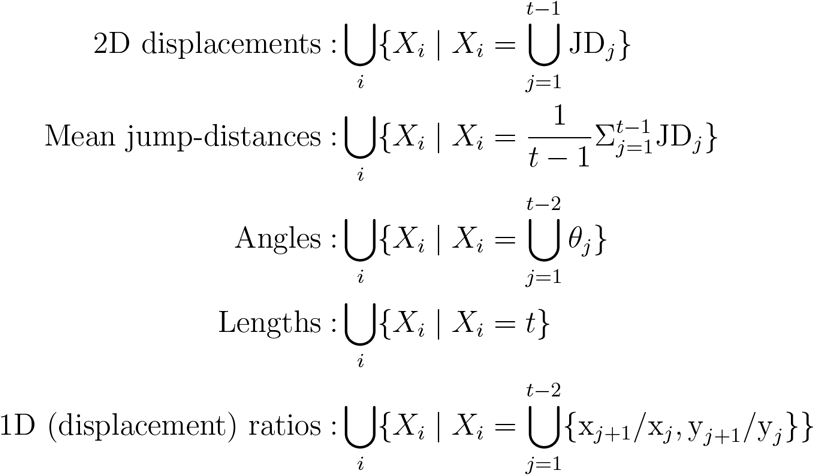

where *X*_*i*_ corresponds to the set of each variable for *i*-th trajectory.

## 2 Introduction of FreeTrace

FreeTrace produces molecular trajectories from a microscopy video, assuming that the trajectories follow fractional Brownian motion. It has a memory for each individual trajectory; hereafter, the trajectory is referred to as a path, which supports the selection of future paths based on the memory of past paths. Unlike Brownian particles, many molecules in the real world do not diffuse independently over time, and their future paths depend on the past paths, such as a dependent stochastic process: fBm (fractional Brownian motion). This memory is the first characteristic of FreeTrace, and the second is the reconstruction of potential paths with multiple time scales. If a particle is detected at time *t* having its past trajectory, FreeTrace explores potential paths until {*t* + 1, …, *t* + Δ*T*} to choose the best among many possible choices. The motivation for this selection on multiple time scales is due to the defocalisation (*e*.*g*., the intensity of molecules, especially immobile ones, fluctuates near the detection threshold) of molecules in real video. Imagine a low-focused molecule; the signal of this molecule is weak, and it alters its acceptance and rejection of detections over time. This happens in every 3D system unless we detect molecules using a threshold (the choice is always binary). In a 2D simulation, this effect is less observable or almost imperceptible, but it is very common in real data. The selection of a path that only considers the interval between *t* and *t* + 1 cannot be free from this limitation in real data and produces an erroneous path, especially for two molecules that travel closely to each other. The selection of a path on multiple time scales seems impossible at first glance since the comparison of probabilities between the paths of different lengths cannot be done in the same probabilistic space. Even if we calculate the analytical density or probability function considering the change of variance depending on time and *H* (Hurst exponent), the obtained analytical density for time Δ*t* is the sum of every potential path reaching the detected particle at *t* + Δ*t* from the particle at *t*. To handle this problem, we treat the potential paths hierarchically, depending on two penalisation terms: one is a penalty for an abnormal jump, and the other is a penalty for a time jump (time gap). By introducing two penalisation terms, paths with the same number of time-jumps and/or abnormal jumps are compared at the same level. In this sense, the selection of the best path (*i*.*e*., the path with the lowest cost) is calculated with the geometric mean of the densities of the path plus its penalisation. The path with the lowest cost is chosen among potential paths. The aforementioned two characteristics are the primary features of FreeTrace compared to other SMT software, and we explain the method in this section, including the relevant mathematical derivations of the density function in fBm.

FreeTrace consists of two major parts: the first is the detection of particles (molecules) from video, and the second is the reconnection of detected particles. We assume the PSF (Point Spread Function) of the measurement instrument is a Gaussian-like function for the detection of particles, and MLE (Maximum Likelihood Estimation) is introduced to find Gaussian-like signals from an image in the section 3. We subsequently present an approximation to estimate the parameters of a Gaussian signal, including the mean, variance, and correlation coefficient of normalised and pixelised photon numbers. The section 4 explains how to reconnect the detected particles under fBm (fractional Brownian motion) on multiple time scales. At the beginning of each section, we provide a brief explanation of the method, along with the notations specific to that section. Note that the notations are different in each section.

## 3 Detection of particles

For the detection of particles from an image, we compare the goodness of fit of two likelihood functions ℒ_0_ and ℒ_1_, which correspond to the Gaussian noise, and the Gaussian PSF added with a Gaussian noise, respectively. The proposed likelihood ratio test to detect particles has already been introduced in MTT [1]. The parameters of two likelihood functions are estimated within *w* × *w*, the size of the sliding window. If the goodness of fit of ℒ_1_ is better than that of ℒ_0_ inside a given sliding window, we accept that this sliding window contains a particle showing a Gaussian PSF at the centre of the sliding window; otherwise, we reject it. For the decision criteria, we use AIC (Akaike Information Criterion). In both ℒ_0_ and ℒ_1_, we assume that the i.i.d. Gaussian noise for each pixel of the image and that the *w* is always an odd number.

### 3.1 Absence of particle inside a sliding window: L_0_

Suppose a sliding window contains only Gaussian noise without any signal of a particle. In that case, we can then fit Eq.1 to the observations ***Y*** inside the sliding window to estimate the mean and variance of the noise *N* with MLE. For ℒ_0_, we have two parameters to estimate, the mean and variance of *N*. we then calculate the partial derivatives of ln ℒ_0_ with respect to the mean and variance. Eq.5 gives the optimal mean 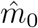, which corresponds to the empirical mean of observations inside the sliding window. Subsequently, Eq.6 also gives the optimal biased variance for the Gaussian noise *N*. We ignored the bias of variance in this step, as the length *w* of the sliding window is typically greater than 7, ensuring sufficient observations, which corresponds to 49 observations inside a sliding window.

**Supplementary Table 2:**
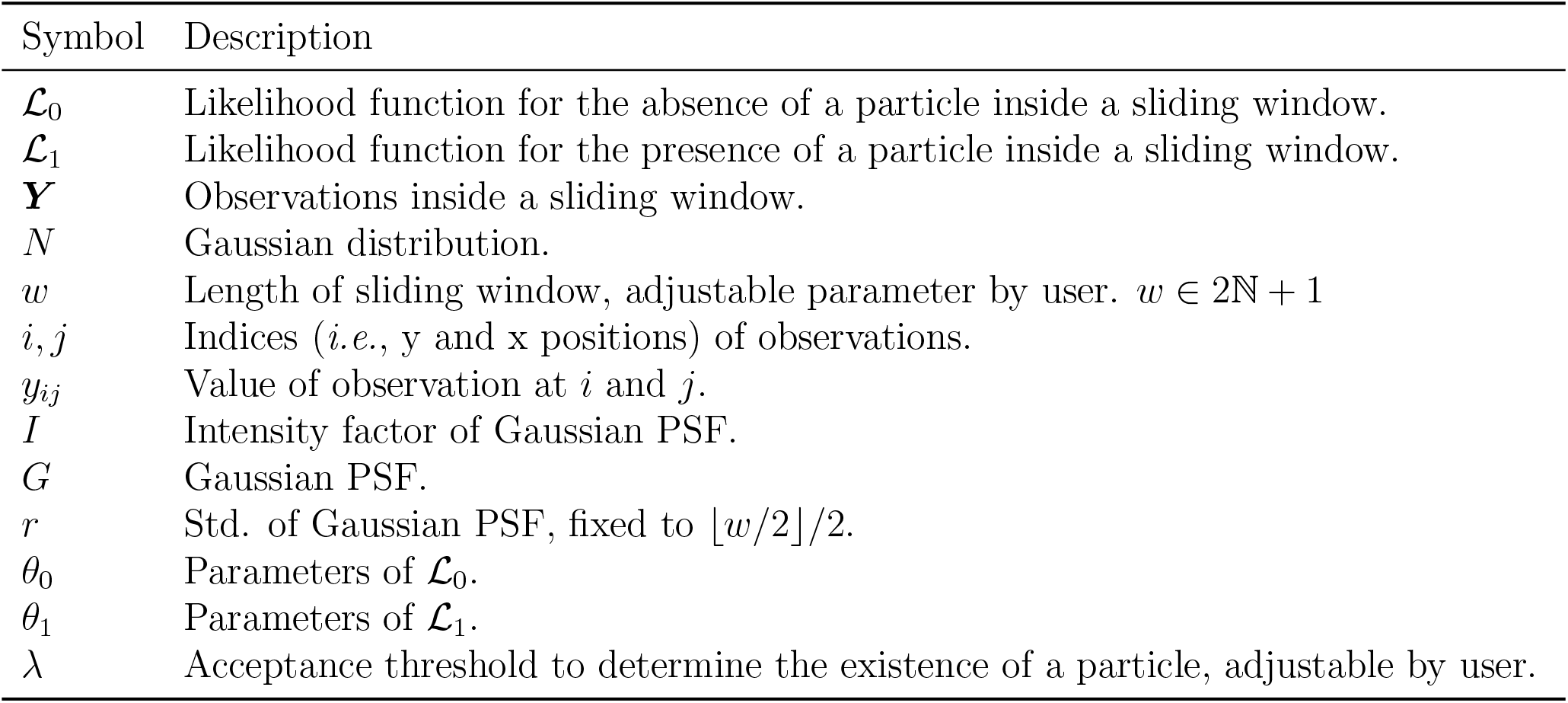
Notation table for the section 3.

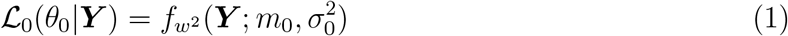

Where 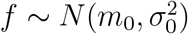. Then, the likelihood ℒ_0_ is

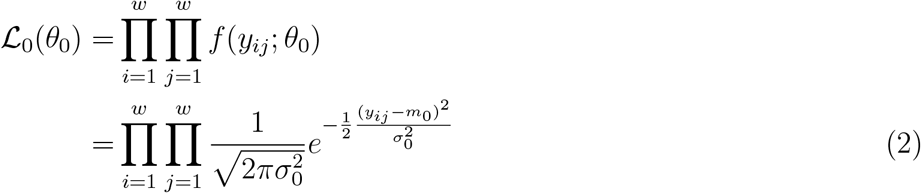

by taking logarithms on both sides,

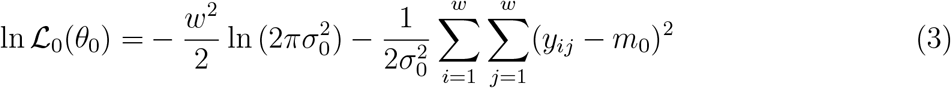

we are looking for *θ*_0_ maximizing the likelihood,

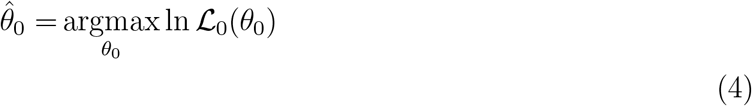

So, the partial derivatives of ln ℒ_0_(*θ*_0_) w.r.t *θ*_0_ are

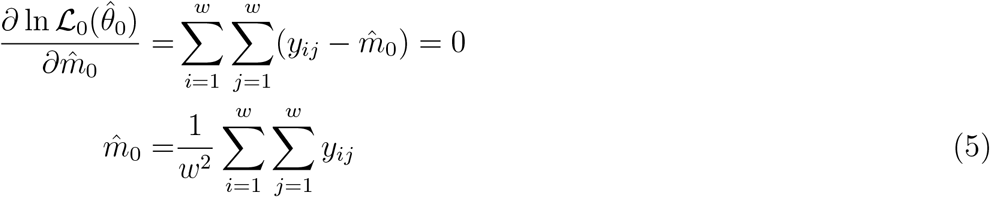

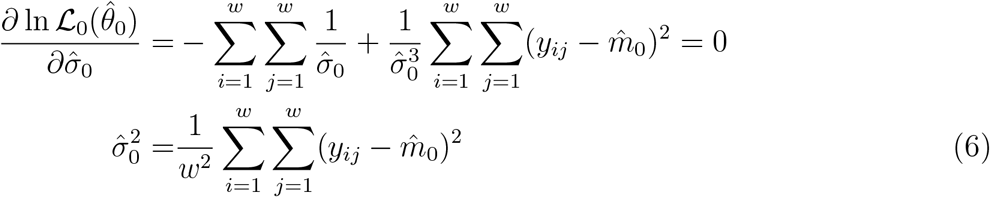

Eq.5 shows that the optimal mean of *N* is the average of observations, and Eq.6 corresponds to the optimal biased variance of Gaussian noise inside a sliding window. The bias correction may produce a meaningful difference for a small sliding window. The 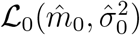 represents the maximized likelihood for the observations ***Y***.

### 3.2 Presence of particle inside a sliding window: ℒ_1_

If there is a particle at the centre of a sliding window with a Gaussian PSF *G* with an additive Gaussian noise *N*, we can fit Eq.7 to the observations inside the sliding window. We fixed that the Gaussian PSF has a variance *r*^2^, where the *r* corresponds to ⌊*w/*2⌋*/*2. We have 3 parameters to estimate in ℒ_1_, which are the mean, variance of *N* and the intensity factor *I* of the Gaussian PSF. Note that we did not estimate the *r* of *G*, because ℒ_1_ can possibly overfit to the very steep signal at the centre of the sliding window. For this reason, *r* is fixed with respect to the length of the sliding window. We calculate next the partial derivatives of ℒ_1_ with respect to the *m*_1_, 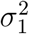 and *I*. Eq.10 and Eq.11 describe the optima for a given fixed *r*, with the estimated optimum of *Î*. In case of *Î*, Eq.12 gives us the optimum, and it corresponds to the sum of the ratio of the observations subtracted by the empirical mean to the Gaussian PSF subtracted by the mean of the Gaussian PSF. The 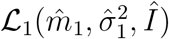 gives the maximised likelihood of the signal of a particle inside the sliding window with Gaussian noise *N*.

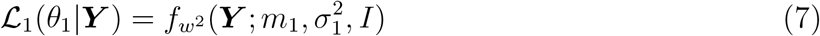

Where 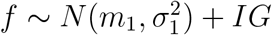.

*G*_*ij*_(*r*) corresponds to the value of Gaussian PSF on 2D plane with fixed *r* for a given position i and j, where *G*_*ij*_(*r*) = 1*/*(2*πr*^2^) exp(−((*i* − ⌊*w/*2⌋)^2^ + (*j* − ⌊*w/*2⌋)^2^)*/*2*r*^2^). Then, the likelihood ℒ_1_ is

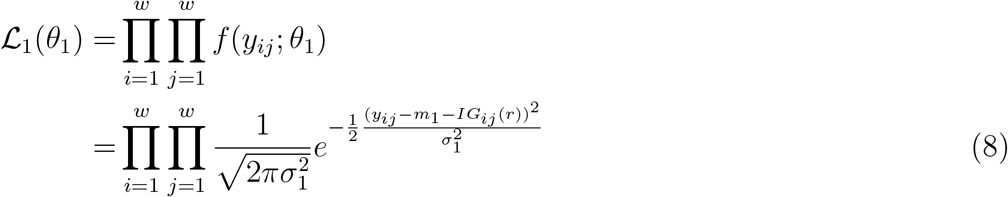

by taking logarithms on both sides,

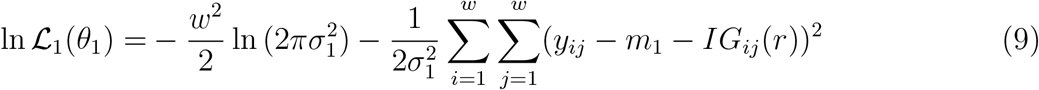

The partial derivatives of ln ℒ_1_(*θ*_1_) w.r.t *θ*_1_ are

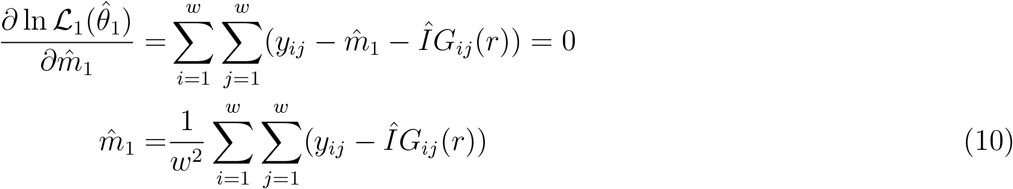

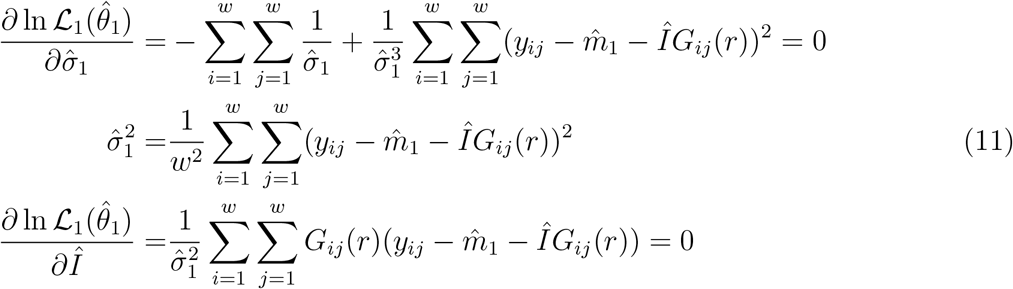

substituting 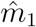 from Eq.10 and letting 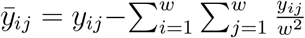 and 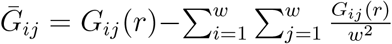,

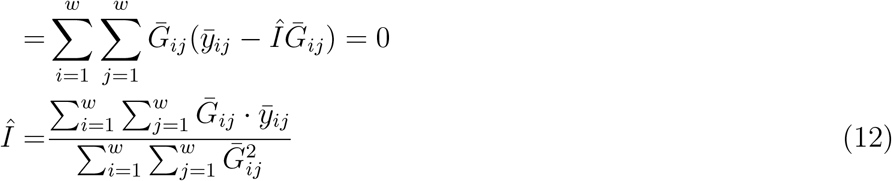

We can obtain the maximized likelihoods of ln 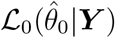 and ln 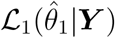 with a fixed *w*.

### 3.3 Decision with AIC

From the maximised likelihood functions ℒ_0_ and ℒ_1_, we compare the goodness of fit for each likelihood from given observations ***Y***. We use the AIC, considering different numbers of estimated parameters for each likelihood in Eq.13.

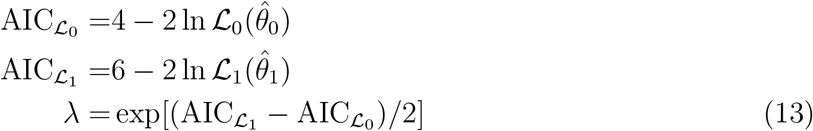

We reject ℒ_1_ if *λ >* 0.05, which represents “we reject ℒ_1_ if the minimized information loss of ℒ_1_ is greater than 95% compared to ℒ_0_”. This corresponds approximately when 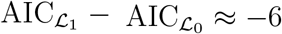. The threshold is adjustable by the user in FreeTrace software.

### 3.4 Gaussian regression to estimate the centre of particle at sub-pixel level

From the detected particle inside the sliding window of size *w*^2^, we can fit a 2D Gaussian function to approximate the position (*x*_0_, *y*_0_) of the particle at sub-pixel accuracy, as well as the x and y variances 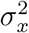 and 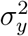, correlation *ρ*, and the intensity factor *I*. We estimate 6 parameters of the Gaussian function as the particle moves constantly and exhibits motion blur, which affects the correlation and each of the variances. Eq.14 minimises the squared loss of the observations ***Y*** to the 2D Gaussian function, which gives the optima of 6 unknown parameters. Assuming the observations are already subtracted by the mean of background noise 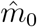, Eq.15 gives the residual *δ* by taking the logarithm on both sides of Eq.14. Note that the residual *δ* is weighted with the observations ***Y***. To solve Eq.17 by minimising *δ*, we need to differentiate *δ*^2^ with respect to 6 unknowns to find the optimum of each. Note that the 6 unknowns of the 2D Gaussian function are converted into a,b,c,d,e and f from Eq.16 to Eq.17 for the simplicity of formulation. The derivatives of *δ*^2^ with respect to each unknown give an overdetermined system as shown in Eq.19. In cases where the empirical distribution shows a heavy-tailed distribution due to signals close to each other, we can iterate the system, which typically ends within 3-4 iterations. The iterative matrix equation to approximate the unknowns is shown in Eq.20. Note that the following approximation for a 1D Gaussian function is already presented by Guo [2], and we employed his approach in a 2D Gaussian function to localise the molecules at sub-pixel accuracy.

### 3.5 Bivariate Gaussian regression of detected particle

The objective function that we want to minimise is as follows:

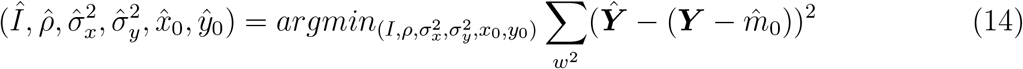

The Above system can be transformed into a weighted linear least squares, and the residual vector *δ* becomes

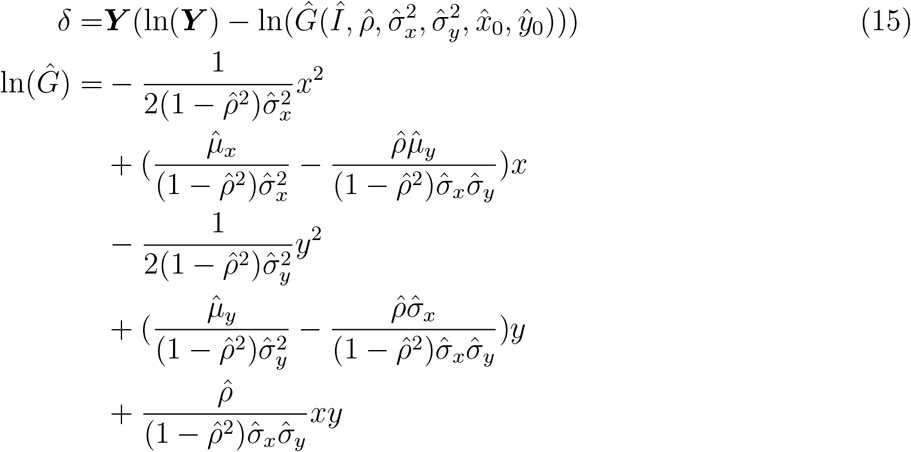

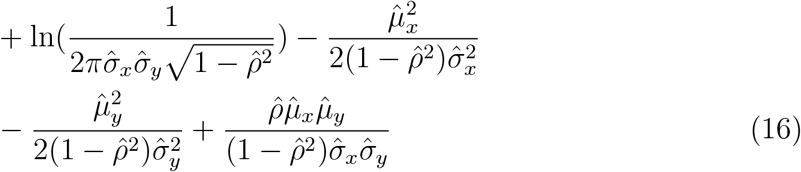

Then, Eq.16 can be converted into Eq.17 as following.

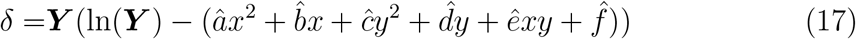

The derivative of *δ*^2^ with respect to *â* to approximate the optimum for 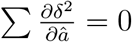 is

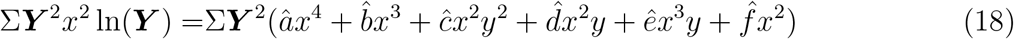

and also differentiate *δ*^2^ w.r.t. 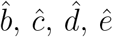 and 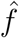, which gives a weighted linear matrix equation for the approximation of 6 unknowns.

### 3.6 Approximate unknowns of determined linear matrix equation

From the Eq.18, we can constructing the following matrix equation,

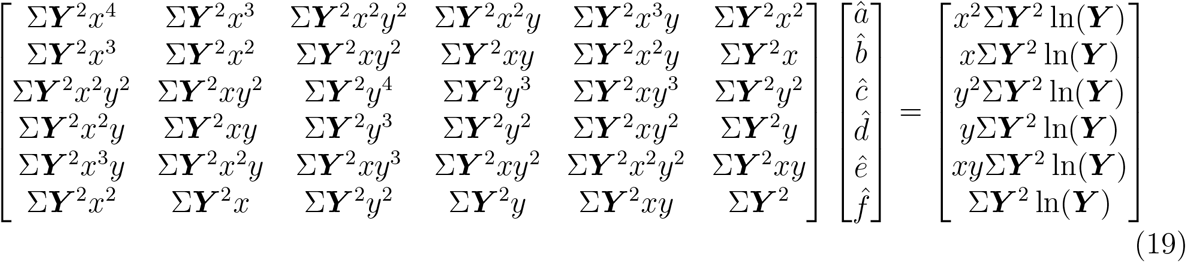

The unknowns of the above linear matrix equation can be approximated with the iterations of the system until convergence. Let *M*_(*k*)_ the matrix at *k*^*th*^ iteration:

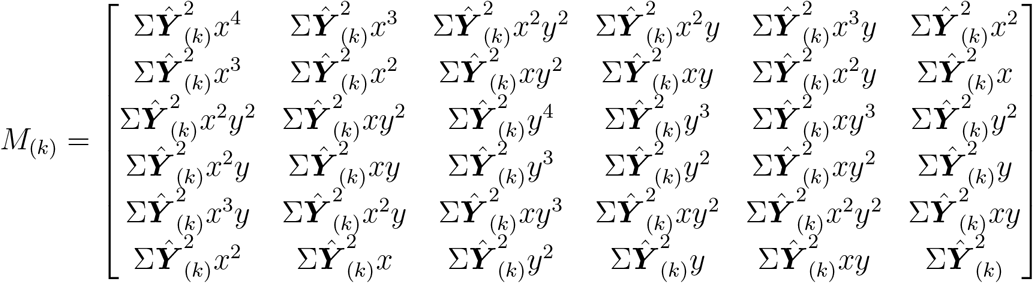

then, the iterative matrix equation becomes

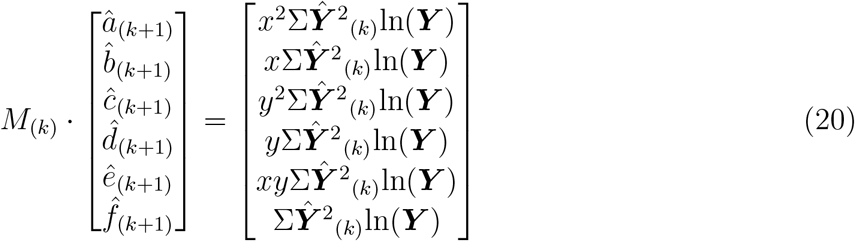

where ***Ŷ***_(0)_ = ***Y***. Finally, we can approximate 6 unknowns of the bivariate Gaussian distribution by solving the above equations iteratively, and the shape of the distribution can be approximated as follows:

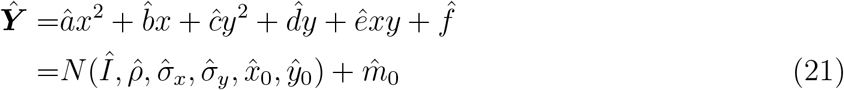

Thus far, we’ve detected and approximated the position of the particle at sub-pixel accuracy within a given sliding window, using two fixed parameters: *w* and *λ*. Note that the methods in the section 3 can be parallelised and detect the particles from a video in real-time.

## 4 Reconnection of detected particles

From the detected particles over time, we reconnect the detected particles to reconstruct trajectories, which are the temporal sequences. Note that the notations in this section (**3**) are different from the section 3.

The reconnections are performed under the assumption of fBm, and we consider that the paths can have different time lags between observations due to blinking events^1^.

In the section 4.1, we describe an algorithm to construct the potential paths from *t* to *t* +Δ*T*. The mathematical derivations are presented in the section 4.2 describing how the ratio of two fGn (fractional Gaussian noise, *i*.*e*., increments of fBm) variances with Δ*s* and Δ*t* can be derived from the standard 1D fBm. The section 4.4 shows an algorithm to select the best path from potential paths.

The reconnection across multiple time scales implies that the selection of the best path in FreeTrace is based on comparisons across multiple frames, not only between the two consecutive time points *t* and *t*+1, to avoid trajectory fragmentation due to blinking. This approach enables us to reduce mis-reconnections (*i*.*e*., interconnections between two distinct paths) caused by blinking events and to prevent overestimation of the number of trajectories from video. We can imagine scenarios where particles are blinking, two particles overpass each other, or the particles go out of the observable space (*i*.*e*., focal depth), but re-enter after Δ*T*. The last scenario occurs especially frequently in real data due to 3D movements within a very narrow observable z-axis range. The event in which particles exit the observable space and re-enter after some time; we’ll call this defocalization hereafter. This defocalization poses a significant challenge to robustly reconstructing molecular trajectories from real data. There are two problems related to defocalization when considering only the two consecutive time points *t* and *t* + 1, rather than multiple frames. The first problem arises frequently when two immobile-like particles are observed closely, and their inter-distance is similar to the commonly observed jump distance of free-diffusive particles. In this case, they can be interconnected due to defocalization of each, if we only consider *t* and *t* + 1, and generate undesirable noises that cannot be observed in jump-distance distributions. If we ignore this problem, the trajectories of these two molecules appear to have no issues with jump distance, since this false jump distance can be observed in free particles; however, they are clearly mis-reconnected. If we analyse the observations subsequently to check whether they change the type of motion, we may mistakenly conclude that their motions change frequently, even though they do not. To reduce these unexpected reconnections, a particle’s memory of its state is needed to avoid interconnections between two distinct particles, even if their inter-distance is probable. The second problem is related to the unexpected fragmentation of trajectories. The unexpected fragmentation of trajectories caused by defocalization reduces the length of the overall trajectories. As a consequence, we underestimate the length of trajectories, which increases uncertainties in subsequent analysis for each individual trajectory due to decreased length. How to reduce this defocalization throughout the overall steps is an important factor in reconstructing and analysing the trajectories in experimental data in SMT.

### 4.1 Reconstruction of potential paths

The potential paths are generated by building a sub-graph **G**_*sub*_, which consists of nodes (*i*.*e*., detected particles in a range of time) and edges (*i*.*e*., paths over time reconnecting detected particles). Δ*s* and Δ*t* denotes two time lags of three observed time points, *t* − Δ*s, t* and *t* +Δ*t*. From a given Δ*T*, which is a fixed parameter denoting the amount of time range to predict at each step, Algorithm 1 accumulates the potential paths for a particle at the time point *t*. Then, we merge the generated **G**_*sub*_ of each particle and return a graph **G** which contains all potential paths from *t* to *t* + Δ*t*. We now have potential paths in **G** from *t* to Δ*t*. The subsequent problem is to choose the most probable paths and determine the number of particles under the assumption of fBm. However, the paths in **G** start from different nodes, each with a different number of paths, and are interconnected at intermediate steps, ending at different nodes. As a result, the choice of the best path is complex due to the different lengths of paths and the different time lags of edges, since the paths are built on different probabilistic conditions. Note that we reconstructed paths with a given Δ*T*, so the edges can have different time lags. *e*.*g*., an edge linking two nodes from *t* = 1 to *t* = 2 and another edge linking two nodes from *t* = 1 to *t* = 3, which do not share the same starting node, cannot be directly compared due to different conditions, even if we can estimate the density (*i*.*e*., probability) for each edge. Thus, FreeTrace simplifies the comparison of paths by approximating their central tendency using the geometric mean of the densities. *e*.*g*., how much the edges of a path are centralised with the analytical expectation of fGn. The most centralised path is selected, and the nodes of the selected path are eliminated from the **G**. This procedure iterates until **G** becomes empty. Thus, we can consider that **G** is emptied in a way of greedy algorithm by selecting the best path at each iteration. This procedure is shown in Algorithm 2. A remaining question is how to compute the density of each edge in **G**. First, we show the auto-covariance of fGn for two time lags Δ*s* and Δ*t*. The analytical auto-covariance of fGn leads to the Cauchy distribution, which is the ratio of two edges (*i*.*e*., the ratio of two 1D displacements). We estimate the density of the ratio from two consecutive edges in a path with the estimated *Ĥ* of the previous path. Since the scale and location parameters are dependent on *H* in the Cauchy distribution, the accurate estimation of *H* for short individual paths determines the overall quality of the trajectory reconstruction result. For this, we estimate *H* using a deep learning model, denoted as *M*. Note that the newborn paths with fewer than 5 observations are assumed to be a Brownian particle with *H* = 0.5 due to high bias and low precision of *H* estimation at the individual level. We show the analytical covariance of fGn in the section 4.2 and the estimation of density for the ratio of two edges in the section 4.3.

**Supplementary Table 3:** Notation table for the section 4.

| Symbol | Description |
| --- | --- |
| $H$ | Hurst exponent. |
| $B^H$ | FBm process with Hurst exponent $H$. |
| $\Delta T$ | Given amount of time (*i.e.*, frames) to predict at each step, fixed parameter. |
| $\Delta t$ | Time lag between the observation at time $t$ and the future observation $t + \Delta t$. |
| $\Delta s$ | Time lag between the observation at time $t$ and the past observation $t + \Delta s$. |
| $\sigma^2$ | Variance of standard fGn. |
| $\sigma_{\Delta t}^2$ | Variance of fGn with $\Delta t$. |
| $\sigma_{\Delta s}^2$ | Variance of fGn with $\Delta s$. |
| $\rho$ | Correlation coefficient of fGn. |
| $\mathbf{G}$ | Graph containing all potential paths of every particle from $t$ to $\Delta T$. |
| $\mathbf{G}_{\text{sub}}$ | Sub-graph containing all potential paths of one particle from $t$ to $\Delta T$. |
| $\mathbf{G}_f$ | Graph containing the selected paths, result of Algorithm 2. |
| $\mathbf{P}_t$ | Set of particles detected at $t^{\text{th}}$ frame. |
| $p_i^t$ | Coordinates of $i^{\text{th}}$ particle at $t^{\text{th}}$ frame. |
| $\text{edge}(a, b)$ | Edge reconnecting two nodes (*i.e.*, particles) $a$ and $b$. |
| $d(a, b)$ | Euclidean distance (*i.e.*, jump-distance) between the nodes $a$ and $b$. |
| $\text{JD}_{\max}$ | Maximum jump-distance of entire particles. |
| $\epsilon$ | Penalty for the path containing a non-temporal-consecutive edge. |
| $\xi$ | Penalty for the path containing an abnormal edge. |

Given coordinates of detected particle, Δ*T* and JD_*max*_, we can reconstruct a **G**_**sub**_ containing potential paths of a particle from *t* to *t* + Δ*T*. Algorithm 1 produces the potential paths of a given particle by accumulating the paths for each frame. The result graph of Algorithm 1 is a multi-DAG (Directed Acyclic Graph). Note that **G**_**tmp**_ is used to avoid the modification of **G** during the iterations for computational reasons.

#### Algorithm 1

Accumulate potential paths for a particle: APP

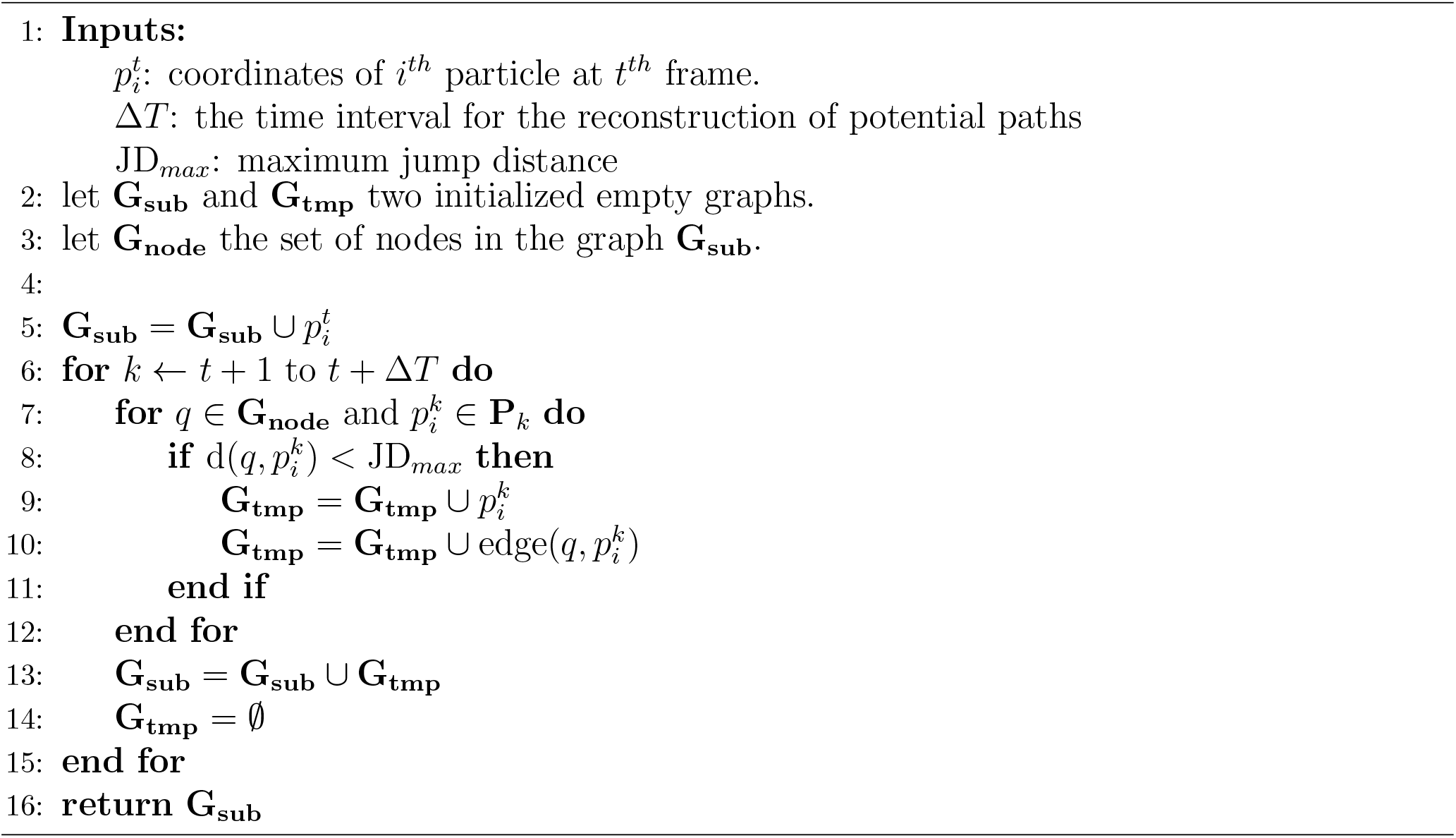

### 4.2 Ratio between two fractional Gaussian noises with different time lags

Supposing that we have 3 observations of a particle under fBm in 1 dimension at the frame *t* −Δ*s, t* and *t* +Δ*t* respectively with the increments 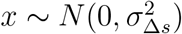 and 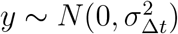. Ideally, Δ*s* and Δ*t* are both 1 everywhere in the 2D simulation since there is no loss of observations due to defocalization. However, they can be different and non-consecutive. This is frequently observed in real data due to the aforementioned reasons, which leads to a difference in variances in fGn. We will see how the autocovariance of fGn can be derived [3, 4] from fBm with different time lags in this section. Let 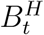 1-dimensional standard fBm, then the increments *X* of 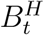 are non-independent and stationary. The increments follow a zero-mean Gaussian distribution *X N* (0, *σ*^2^), where *σ*^2^ is the variance of fGn with a given time span. The covariance of two fBm for two arbitrary time points *n* and *m* can be calculated as follows using the property of self-similarity, assuming *m > n*,

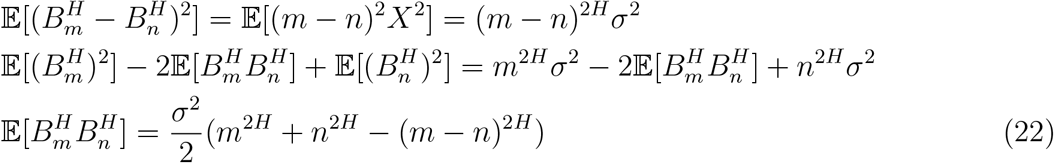

The Eq.22 is already well-known autocovariance function of fBm with respect to *H*, and *σ* corresponds to 1 for standard fBm. However, we will focus on the autocovariance of fGn instead of fBm. The autocovariance of two fGn with two different time lags Δ*s* and Δ*t* can be shown as follows:

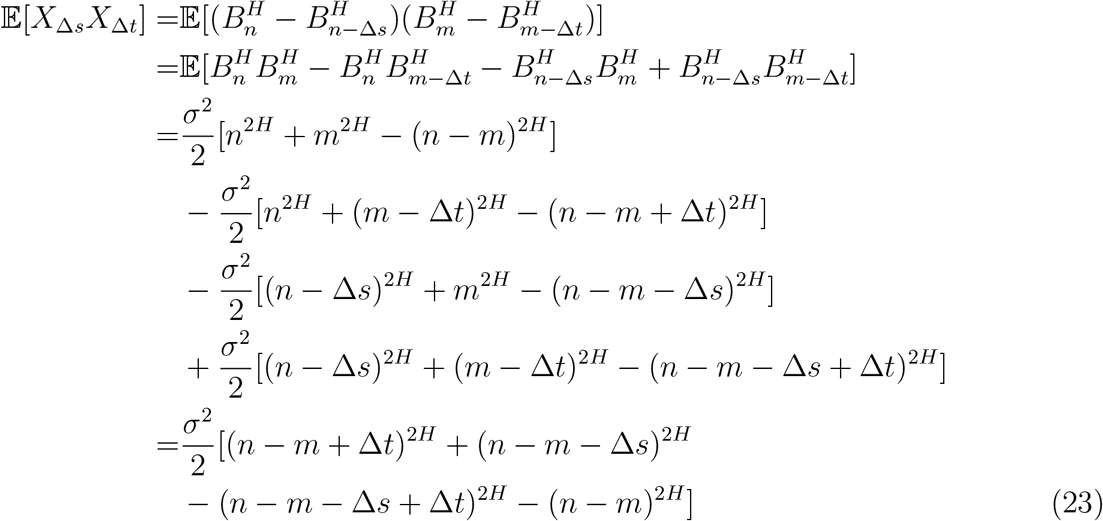

we are interested in the autocovariance of fGn when *m − n* = Δ*t*, Thus, Eq.23 becomes

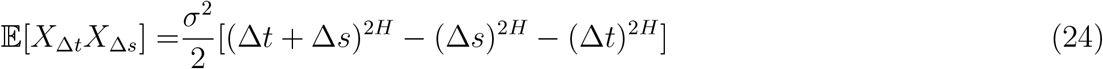

When Δ*t* = Δ*s*, the autocovariance for any time lag Δ*n* of fGn is

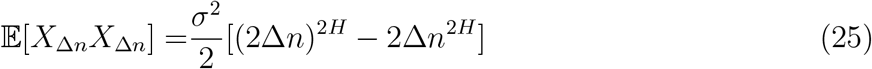

We can obtain the standard correlation coefficient *ρ* when Δ*n* = 1,

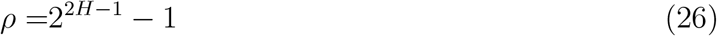

We can also consider the ratio of autocovariances of fGn with respect to different time lags but with the same *H*,

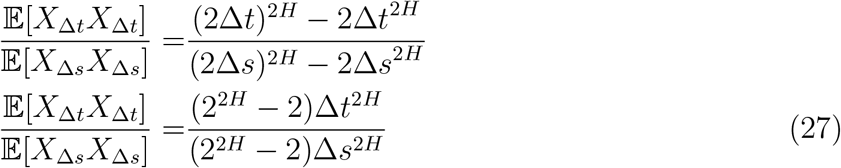

We can see how autocovariance of fGn evolves with respect to time lags and *H* and the ratio with Eq.25 and Eq.27.

Assuming that the time only affects the variance of fGn, not the correlation coefficient,

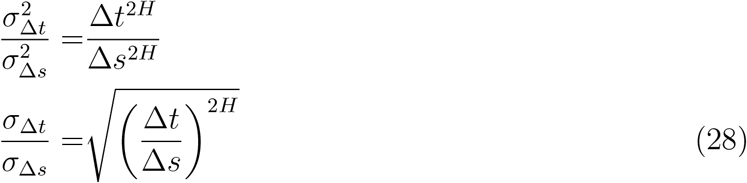

We can see the ratio of the standard deviation of two fGn with different time lags Δ*s* and Δ*t* in the Eq.28.

### 4.3 Density estimation with Cauchy distribution

We showed the ratio of standard deviations in Eq.28 for two observed distances in 1D with different time lags. we can estimate the density of *y/x* where *x* and *y* are two observed and correlated distances. The underlying distribution of this ratio corresponds to the Cauchy distribution since both *x* and *y* follow a zero-mean Gaussian distribution by the definition of fBm. The derivation of the Cauchy distribution for *y/x* can be shown from the standard bivariate Gaussian distribution as follows,

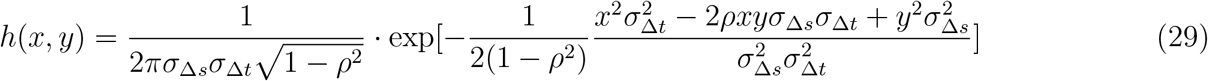

with *u* = *y/x*,

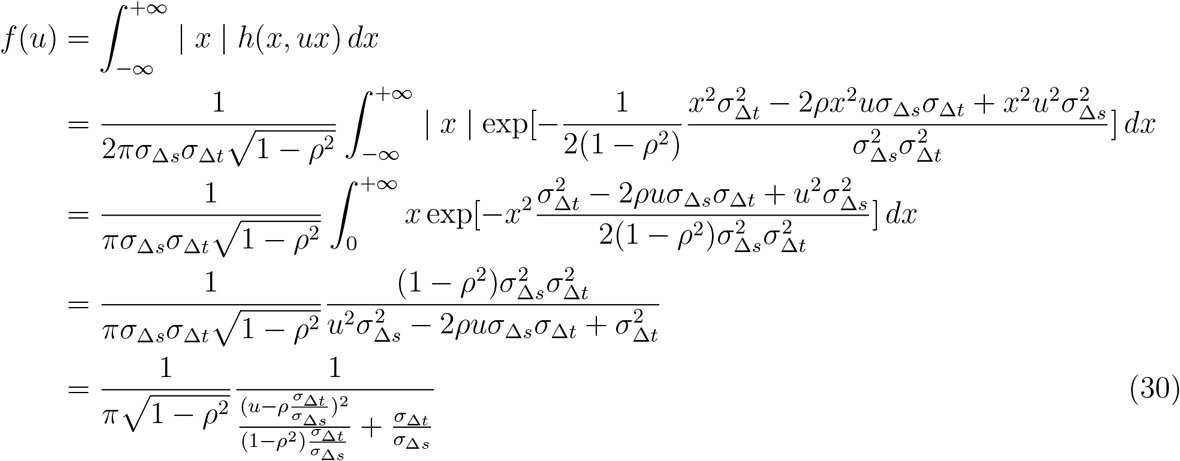

The Eq.30 is Cauchy distribution with a location 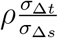.

Then, the density of the displacement-ratio with two different time lags can be obtained as follows, assuming that *H* is already known for the current path,

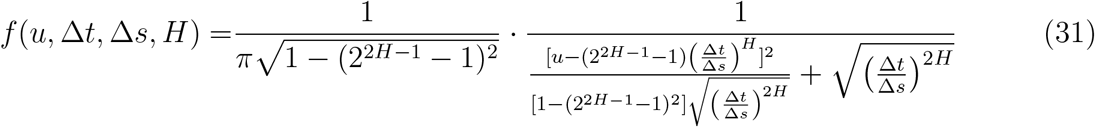

The distribution function of equation 31 is as follows,

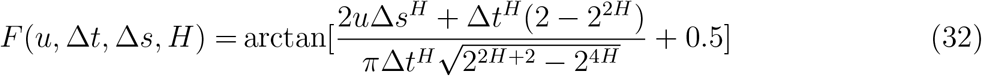

and the quantile function is as follows,

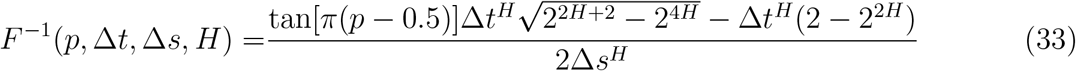

### 4.4 Selection of potential trajectories

We compare the geometric means of path densities to select the best path using a Cauchy distribution. Since **G** is a multi-directed acyclic graph (DAG), finding the global optimal path is NP-hard, as with TSP (Travelling salesman problem). Thus, we find the best path greedily in such ways. From a given **G**, we select first the highest-density path and exclude the selected nodes from the **G**. At the second iteration, we select again the highest-density path from **G**. This iteration continues until **G** becomes empty. This strategy doesn’t guarantee the global optimum. However, it can find the sub-optimal **G**_*f*_ in reasonable computation time when the size of **G** gets larger. Note that the path, including the edge linking two nodes that are non-consecutive in time and the edge linking two nodes where the ratio is outside the *n*-scale range in the Cauchy distribution, is penalised. For example, a path containing an edge between *t* and *t* + 3 is penalised with 2 *· ϵ* and an edge with | 𝔼 (*u*) − *u* | *> n·γ* is penalised with *ξ* where *u* and *γ* are the ratio and scale, respectively. Algorithm 2 and 3 show how the value of the path (*i*.*e*., geometric mean of densities with penalisation) is calculated. In summary, a path has the highest value when the jump-distance ratios are centralised at most from the Cauchy distribution without penalisation. Note that the density of the first edge is computed with the sampled empirical jump distribution since the first edge cannot have a ratio. The *H* value for each path is estimated using a deep neural network (ConvLSTM) [5] pre-trained with simulated fBm paths.

From Eq.31, we can calculate the density of a pair of two consecutive edges with *Ĥ*, where *Ĥ* is the estimated Hurst exponent from the previous path of the current edge. If it doesn’t have the previous path, we consider *H* as 0.5, which means that the particle follows classical Brownian motion. This allows us to choose the next particle depending on the state of the particle’s previous path, and the choice of selection is decided by the memory. The computation of densities from a given path is shown in Algorithm 3 and the selection of the best path from **G** is shown in Algorithm 2. Since the estimation of the first edge for each path cannot be estimated with a ratio, we estimate it with the empirical jump-distribution *g*(*y*, Δ*t*) which gives the empirical density of *y* for Δ*t*.

## 5 The depth of the graph for trajectory reconstruction

The depth of the DAG, Δ*T*, is an important feature for reconstructing trajectories, which defines the number of possible paths considered within a time span. To identify the appropriate number of Δ*T* on the simulated scenarios, we tested FreeTrace on the low density of the scenario of **Fig. 3a**. The averages of DTW distances (**Supplementary Fig. 3a**) with respect to Δ*T* indicate FreeTrace performs at best when the depth of the graph is equal to three. The increased depth reduces the total number of predicted trajectories as a gap frame in other software. As a consequence, the high number of Δ*T* doesn’t necessarily increase performance; rather, it increases the computational time, which grows factorially depending on the number of neighbouring molecules.

### 6 Estimation of diffusion properties

#### 6.1 At individual level

The estimation of *H* for each individual trajectory is performed using a neural network (ConvLSTM) [5], which has already been shown to have potential in the proposed method [6]. This showed the accurate estimation of *H*. However, it is also shown that this leads to a high bias for short trajectories near *H* equals to 0 and 1. It is intractable to estimate *H* exactly for short trajectories due to the low number of observations (*e*.*g*., 5-length of sequence). However, the Cauchy distribution presented in Eq. 31 and Eq. 32 can be utilised to estimate *H* at the ensemble level, which is presented in the next section.

##### Algorithm 2

Selection of paths

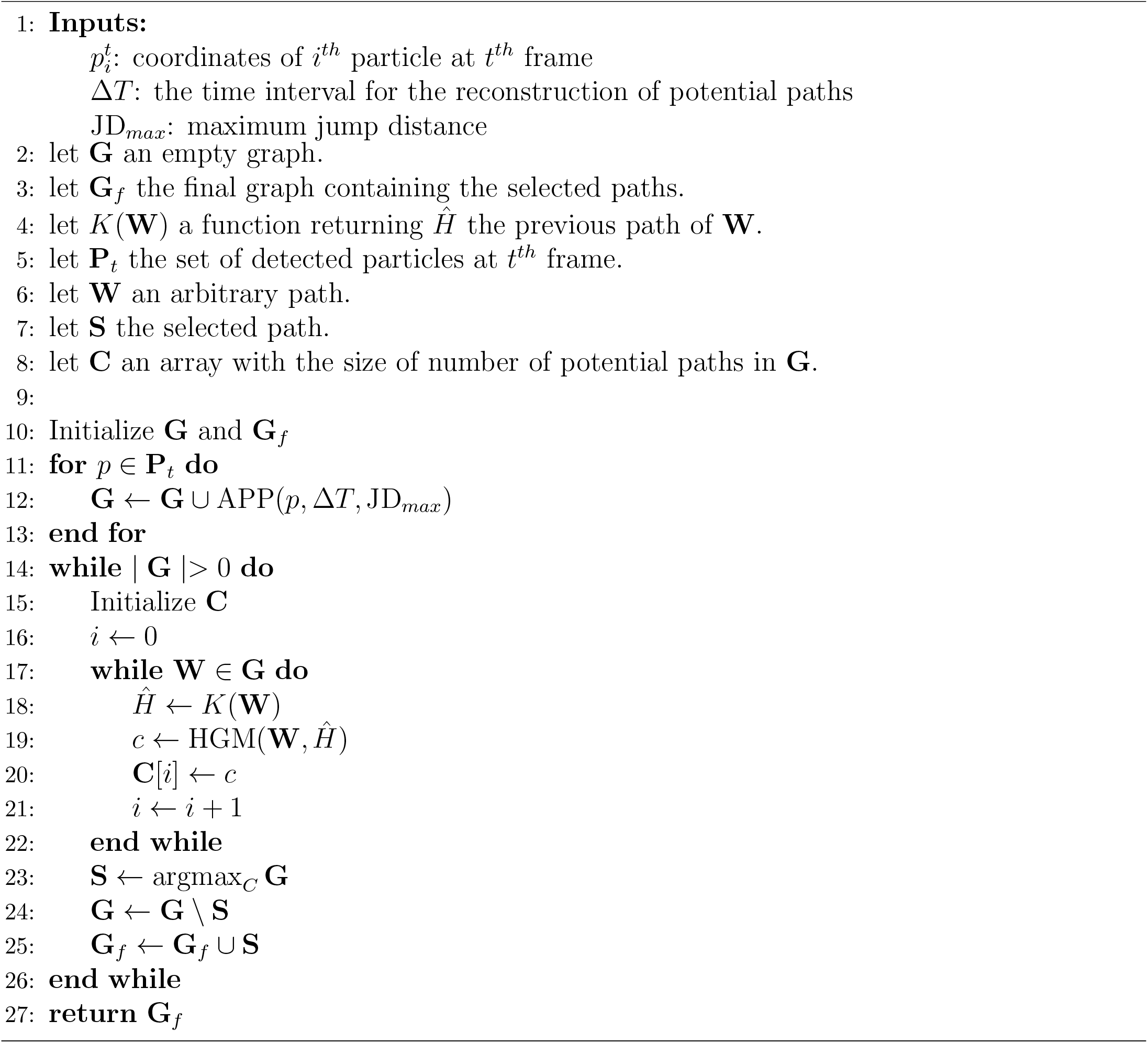

##### Algorithm 3

Hierarchical geometric mean: HGM

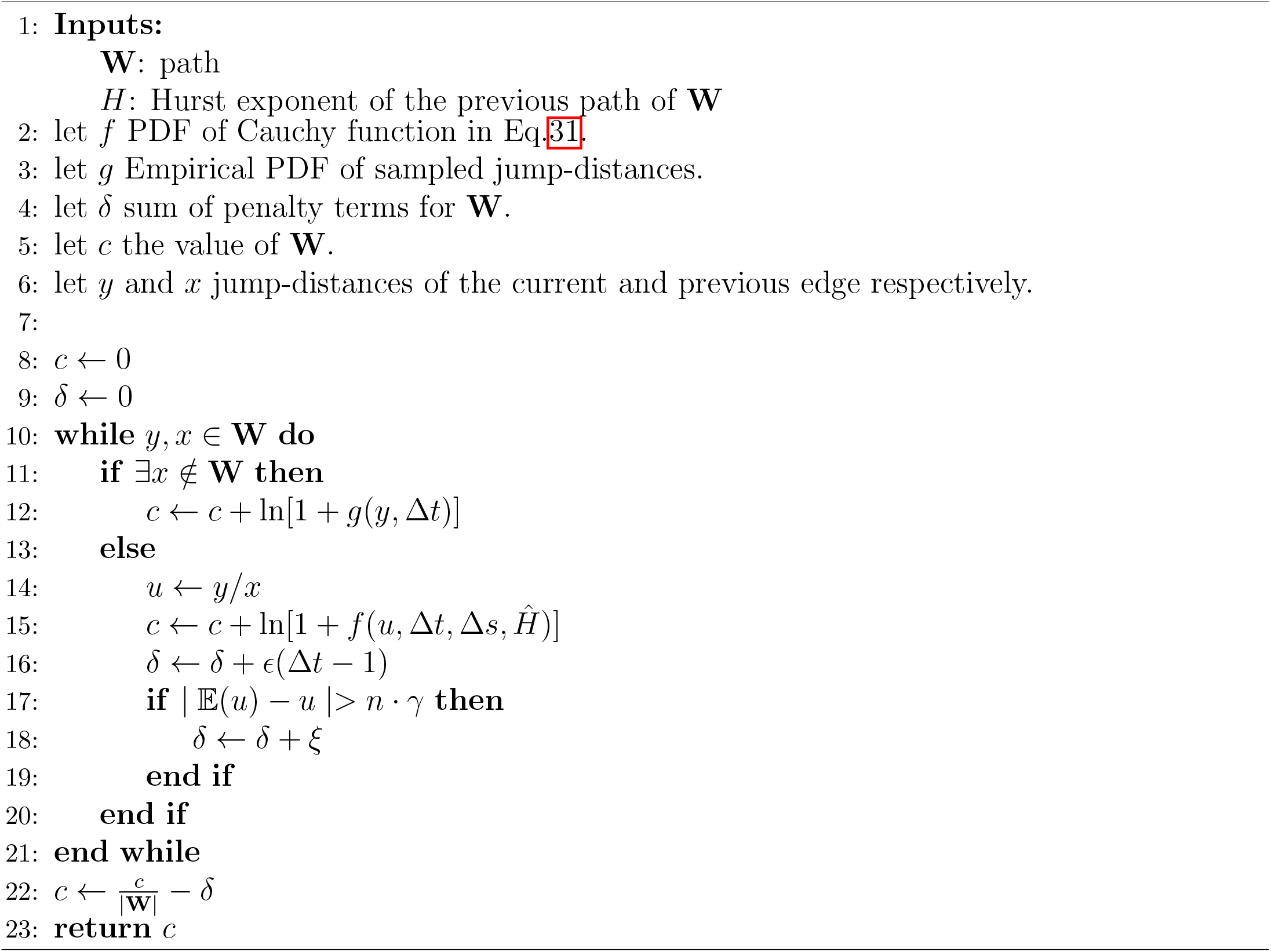

The *K* value for each individual trajectory is estimated using a relatively simple network, as shown in the proposed method [6].

#### 6.2 At ensemble level

Supposing that a trajectory has a constant diffusion coefficient throughout the observation period. Then, the ratio of increments eliminates the diffusion coefficient, and only the temporal correlation of increments remains, as shown in Eq. 31. This signifies that we can fit the Cauchy distribution to estimate the *H* even for short trajectories, assuming that they belong to a single population. For example, if we have 10,000 2-length (3 frames) trajectories belonging to the same population, we can then obtain 10,000 ratios. Estimating *H* and *K* for each individual trajectory is almost impossible in this case. However, we can estimate their *H* at the ensemble level, as the estimation of the diffusion coefficient in Brownian particles, using the proposed Cauchy distribution, regardless of the length of the trajectories. The accuracy in this case is naturally proportional to the number of observations of the total ratios of a population (**Extended Data Fig. 1**). We utilised the non-linear least squares method for the fitting of the Cauchy distribution with truncation, but MLE may be utilised with further investigations.

We did not introduce a method to estimate *K* at the ensemble level throughout the paper, since its estimation is already well-introduced in [7].

## Footnotes

1 the movement of a particle in 3D space distorts the PSF and results in blinking events alternating between detection and non-detection over time

